# Differentiation is accompanied by a progressive loss in transcriptional memory

**DOI:** 10.1101/2022.11.02.514828

**Authors:** Camille Fourneaux, Laëtitia Racine, Catherine Koering, Sébastien Dussurgey, Elodie Vallin, Alice Moussy, Romuald Parmentier, Fanny Brunard, Daniel Stockholm, Laurent Modolo, Franck Picard, Olivier Gandrillon, Andras Paldi, Sandrine Gonin-Giraud

## Abstract

Cell differentiation requires the integration of two opposite processes, a stabilizing cellular memory, especially at the transcriptional scale, and a burst of gene expression variability which follows the differentiation induction. Therefore, the actual capacity of a cell to undergo phenotypic change during a differentiation process relies upon a modification in this balance which favors change-inducing gene expression variability. However, there are no experimental data providing insight on how fast the transcriptomes of identical cells would diverge on the scale of the very first two cell divisions during the differentiation process.

In order to quantitatively address this question, we developed different experimental methods to recover the transcriptomes of related cells, after one and two divisions, while preserving the information about their lineage at the scale of a single cell division. We analyzed the transcriptomes of related cells from two differentiation biological systems (human CD34+ cells and T2EC chicken primary erythrocytic progenitors) using two different single-cell transcriptomics technologies (sc-RT-qPCR and scRNA-seq).

We identified that the gene transcription profiles of differentiating sister-cells are more similar to each-other than to those of non related cells of the same type, sharing the same environment and undergoing similar biological processes. More importantly, we observed greater discrepancies between differentiating sister-cells than between self-renewing sister-cells. Furthermore, a continuous increase in this divergence from first generation to second generation was observed when comparing differentiating cousin-cells to self renewing cousin-cells.

Our results are in favor of a continuous and gradual erasure of transcriptional memory during the differentiation process.

## Introduction

During cell division, the mother-cell endures a period of transient instability – the mitosis – which is accompanied by dramatic cellular and epigenomic reorganizations [1]. The close to equal partitioning of the cellular content, together with active mechanisms, such as the conservation of gene transcription profiles after division by chromatin-related epigenetic mechanisms, or the long half-life of proteins ensure the overall phenotypic similarity of the sibling cells [2–4]. As a consequence, the resulting sister-cells regain immediately after the division many of the structural and functional features of the maternal cell. The phenotypic stability of clonal cell lines is largely founded on this phenomenon frequently called “cellular memory”.

A small number of studies have addressed the question of the preservation of cellular memory through division using different approaches ranging from microfluidics combined with scRNA-seq [5], to time-lapse microscopy of reporter genes expression [6, 7], to a dedicated procedure called MemorySeq [8]. Those studies have been focused on self-renewing cells, such as mouse ES cells or melanoma cell line. In all cases, the authors concluded to the existence of a transcriptional memory defined by the heritability of gene expression levels in a gene-specific manner, extending up to two or more generations. This transcriptional memory impacts subsets of genes called “memory genes”, the expression of which is uncorrelated in a population of cells but correlated in sister-cells. Those genes are highly dependent on the cell system used for the investigation. Beyond their actual function, the fact that related cells harbour correlated expression for those genes is a read-out for this transcriptional memory and demonstrates the existence of a constraint imposed to the cells gene expression profile at division.

On the other hand, all cellular processes are subjected to stochastic molecular fluctuations which will favor the decorrelation of the sister-cells phenotypes and increase the transcriptional heterogeneity in a clonal population of siblings. For example, relaxation experiments demonstrated on various cell systems that after two weeks of culture under stable conditions, the expression level of specific genes in a selected homogeneous cell clone becomes as heterogeneous as it was in the original population the founder cell derived from [9]. Moreover, the capacity of a cell clone to reconstitute the heterogeneity of the original population over time has been observed in many instances in normal or pathological cell types [4, 8, 10].

During the process of differentiation, this whole delicate balance between the two opposing forces of the stabilizing cellular memory and change-inducing gene expression fluctuations has to be somehow revisited. Indeed, differentiating cells undergo substantial morphological and functional changes.

Although differentiation usually takes place over several cell cycles, there is a critical transition period characterized by stochastic gene expression and rapid morphological fluctuations. A large range of experimental studies have indeed demonstrated, that the first step in cell differentiation is the rapid and transient increase of the variability in gene expression in response to the stimuli inducing the differentiation, both *in vitro* [11–19] and *in vivo* [20, 21].

An important unresolved question is therefore to understand how the dynamic stability and the capacity of differentiation are integrated into a single process. In the present study we aimed to investigate the dynamic balance of stability/instability in dividing cells that undergo the first steps of differentiation. To do this, we measured the resemblance of the sister-cells by comparing their transcriptomes.

We formulated 3 hypotheses on the possible evolution of transcriptional memory upon differentiation induction (Figure 1). To illustrate those hypotheses, cells in a self-renewing state are positioned in a gene expression space (grey sphere). Assuming the existence of transcriptional memory in our self-renewing cells after mitosis, like in other cell models, sister-cells start in roughly at the same position in that space (blue family tree). Then, upon differentiation induction (red family tree), we can postulate the following three hypotheses:

**Figure 1:**
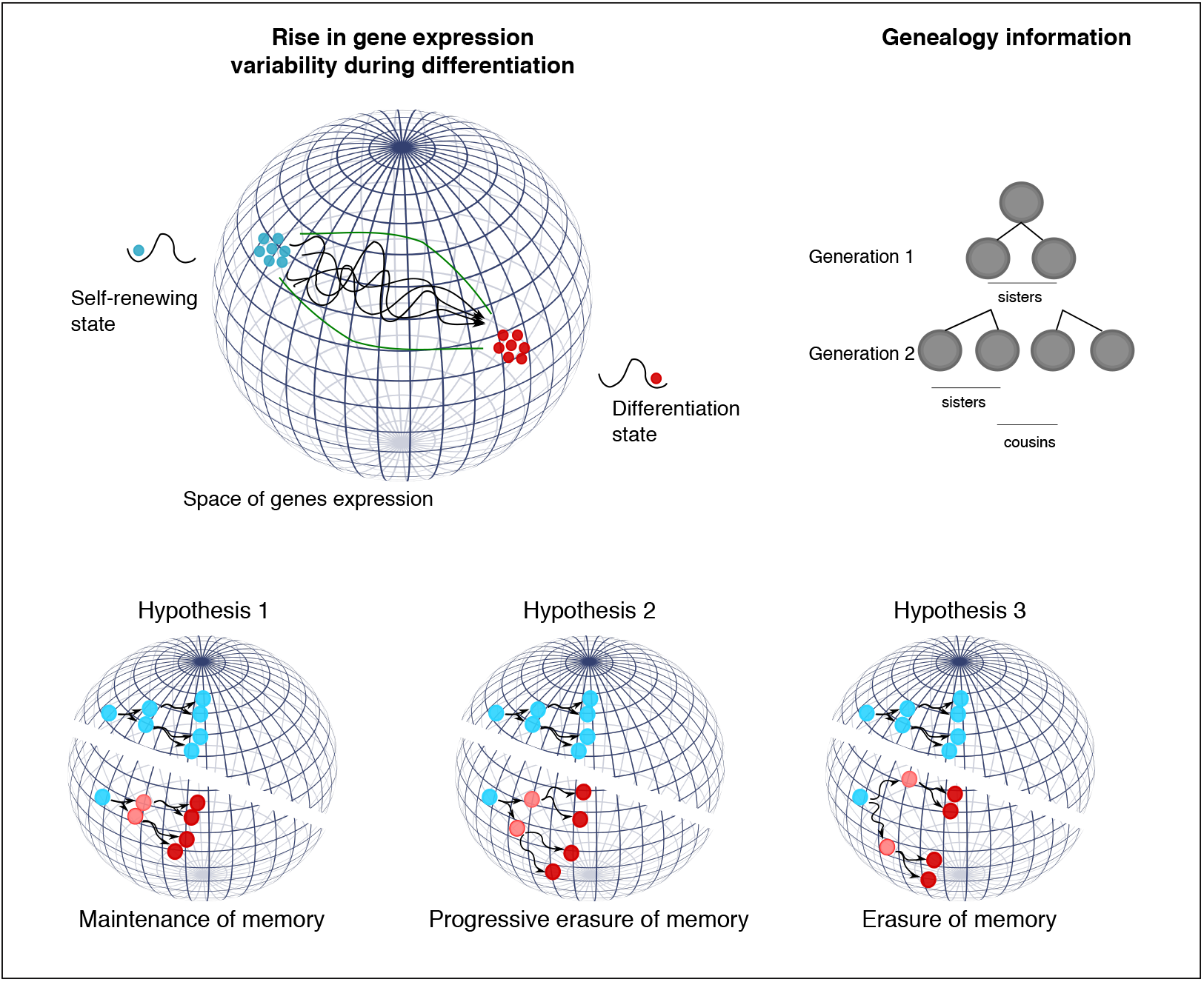
Hypotheses on transcriptional memory during a differentiation process. Self-renewing cells (blue cells) are compared to differentiating cells (red cells) after one and two divisions.

- The maintenance of memory hypothesis: the transcriptional memory overrules the expression variability resulting in related cells following roughly the same path in the gene expression space toward the differentiated state (hypothesis 1), or
- The progressive erasure of memory hypothesis: the memory is gradually erased, translated in our projection to differentiating sister-cells starting to follow roughly the same path and progressively bifurcating from each other, and even more after one more cell division (hypothesis 2), or
- The instantaneous erasure of memory hypothesis: the variability of gene expression pushes the balance and takes over the transcriptional memory, leading each differentiating sister-cell to follow a completely different path from the beginning of the differentiation process (hypothesis 3).

In order to distinguish between those different scenarios, it is necessary to quantitatively evaluate, at the single-cell level, the similarity of the gene expression profiles of sister-cells shortly after the division under self-renewing versus under differentiation-promoting conditions.

Therefore, we developed two strategies to isolate cells while preserving their precise lineage information after one (generation 1) and two (generation 2) divisions, a manual one and a FACS-based one. Then, in order to assess the genericity and robustness of our findings, we compared two different cell differentiation models (human CD34+ cells and T2EC chicken primary erythrocytic progenitors) and for the T2EC model two cellular states: self-renewing and differentiating. We used two different single-cell transcriptomics methods: a highly sensitive targeted quantification method, sc-RT-qPCR and a whole-transcriptome approach, scRNA-seq.

We obtained qualitatively very similar results using the two cell types and the two single-cell measurement technologies. First, after one cell division (generation 1) in both models, and in both states for the T2EC model, we detected a transcriptional memory demonstrated by the sister-cells displaying more transcriptomic similarity between each other than two randomly selected cells. Second, using the T2EC model, which allows to compare sister-cells induce to differentiate to sister-cells in self-renewing state, we also observed that this transcriptome similarity decreased during the differentiation process as compared to the self-renewing cells. Interestingly, this effect was even more pronounced one division later (generation 2), when interrogating cousin-cells. Altogether our results point toward a continuous gradual loss of transcriptional memory during the differentiation sequence.

## Results

### Cellular models of differentiation

To consolidate our results we used two different cell differentiation models. As a first model, we used primary human cord blood derived CD34+ cells. These cells are believed to be a mixture of so-called multipotent progenitors and stem cells that retains the capacity to differentiate into various cell types. Under *ex vivo* conditions, the CD34+ cells, unless stimulated, are stopped in the cell cycle and survive only a few days. When stimulated with a mixture of cytokines, they re-enter the cell cycle and will differentiate into two different committed progenitors [15]. Briefly, by 24hrs after stimulation, a burst in transcription produces a mixed transcription profile called “multilineage primed” state [11] and by the end of the first cell cycle (between 40 and 60hrs), cells with two different transcription profiles emerge in the population [15, 22]. However, this first fate-decision is a highly dynamic and fluctuating process which is more complex than a simple binary switch between 2 options [15]. In the present work, we investigated by sc-RT-qPCR the transcriptional profile of couples of CD34+ sister-cells derived from the first cell division after the cytokines stimulation.

As a second model, we used chicken primary erythrocytic progenitors called T2EC [23]. Contrary to the human cord blood CD34+ cells, these cells can be maintained in a self-renewing state *in vitro* under appropriate culture conditions [24]. They can be induced to differentiate at will into mature erythrocytes by a change of medium [24]. The T2EC cells undergo a simple “switch”: they leave the self-renewing phase and enter a differentiation trajectory without bifurcation toward different end point phenotypes. This model allows a direct comparison of related cells in two different states: self-renewing and during differentiation. Furthermore, a previous study on this model had highlighted a critical point of cell commitment, 24hrs postdifferentiation induction characterized by the rise in gene expression variability, measured with entropy [25]. Thus, we focused on the first steps of T2EC differentiation and investigated the transcriptional profile of couples of generation 1 sister-cells in both cellular states and families of generation 2 sisters and cousin-cells in both state by a scRNA-seq approach [26].

## Cells isolation

### Isolation of first generation cells

We achieved the technical challenge to isolate related cells following their first and second division (generation 1 sister-cells and generation 2 sisters and cousin-cells). The usual molecular tagging or barcoding lineage tracing approaches could not be used in our case since these approaches allow retrieval and analysis of cells belonging to the same clones at later stages, but not at the scale of one cell division [27]. For our investigation, a direct observation of the dividing cells and individual isolation of the generation 1 sister-cells, and generation 2 sisters and cousin-cells were necessary. Furthermore the use of primary cells, with a short life span, precluded the possibility to genetically engineer reporter systems.

We first developed two different methods to recover generation 1 sister-cells, depending upon the cellular model at hand: a manual one and a cytometry-based method. Those original strategies are presented below and in Figure 2. The technical details are explained in the Methods section.

**Figure 2:**
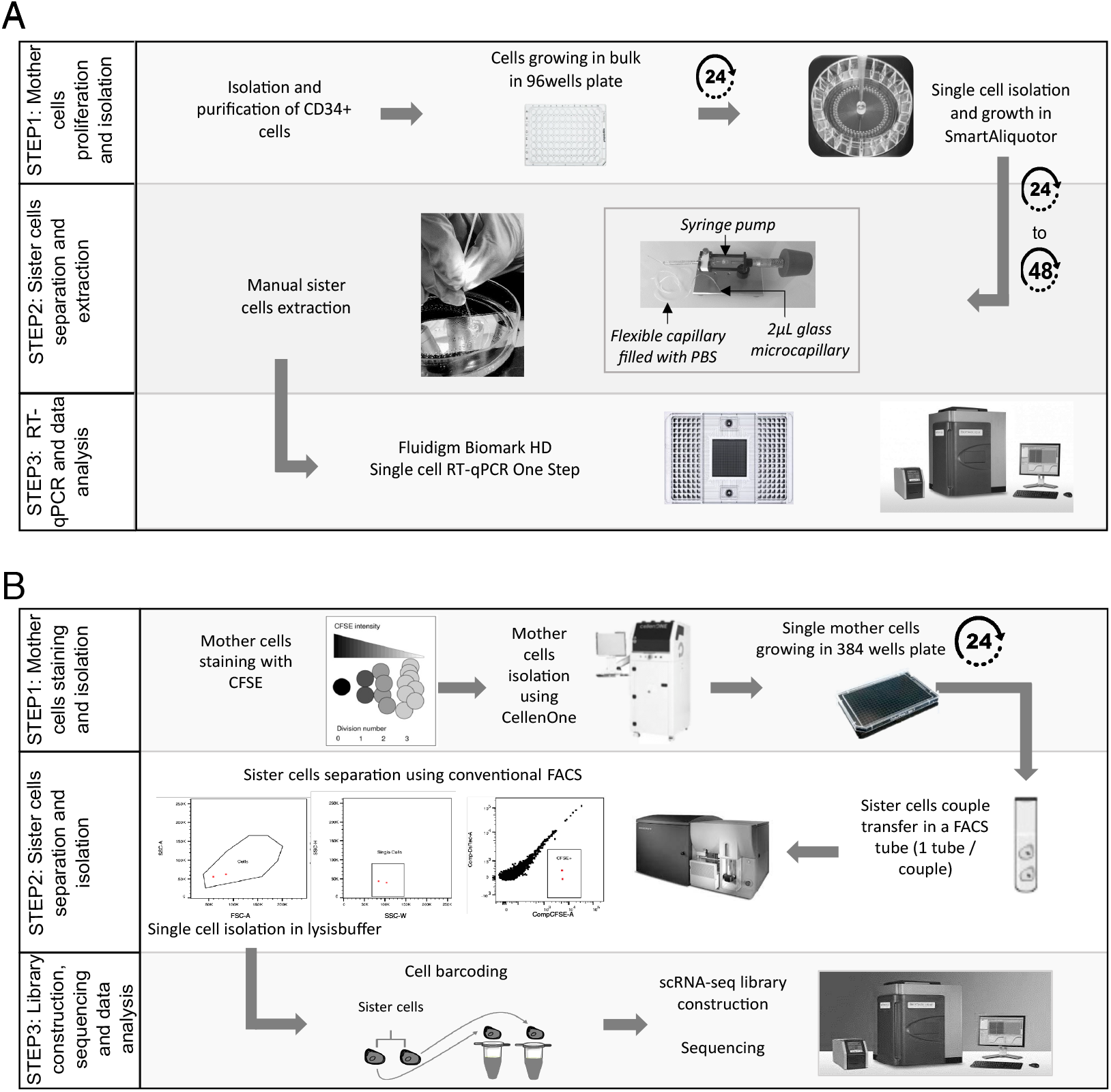
General workflows developed to generate, follow and separate generation 1 sister-cells from CD34+ (A - manual strategy) or T2EC (B - cytometry-based strategy) mother cells. See text and Methods for details.

Human CD34+ cells were grown during 24hrs in a standard 96-well plate before being isolated into single cells, using a Smart Aliquotor device in which individual cells still share the same medium. Isolated mother-cells were then cultured for 24 to 48hrs in the device to allow one cell division. The wells were regularly inspected to detect this first division. Then, the resulting sister-cells were isolated manually under a microscope using a pressure controlled microcapillary and recovered in lysis buffer for further processing. The cells transcriptomes were analyzed by single-cell quantitative RT-PCR using the Fluidigm system as described here [15].

T2EC mother-cells were isolated after CFSE - carboxifluorescein diacetate succinimidyl ester - staining using CellenOne ®low-pressure cell sorter and plated in a 384-well plate. Cell doublets, resulting from the first division, were identified using an inverted microscope. The two cells were then isolated using an FACS Aria cytometer and recovered directly in tubes containing lysis buffer and scRNA-seq primers, for which the cell barcodes sequences were known in advance. scRNA-seq libraries were then constructed as previously described here [28] and sequenced.

Successfully recovering the two sister-cells using FACS is *perse* a remarkable achievement, as this method usually requires hundreds of cells to start with, whereas the initial population here consisted of two cells. To achieve this, we first used the CFSE fluorescence intensity to ensure that the objects isolated were indeed cells (Figure S1 A-B for self-renewing medium and C-D for differentiating medium). CFSE stably binds to the amine groups present in cytoplasmic proteins, conferring stable fluorescence intensity to the cell. As total protein content is supposed to be relatively equally distributed between sister-cells during cell division, so is the fluorescence intensity [29, 30]. We used this specification to validate that the two cells isolated were actually sister-cells. We evaluated the CFSE intensity correlation between pairs of sister-cells, and compared it to intensity correlation values of randomly paired cells from the same dataset (Figure S1 E-F for self-renewing cells and G-H for differentiating cells). Outstandingly, CFSE correlation values between self-renewing sister-cells and differentiating sister-cells were extremely high (0.91 and 0.95 Figure S1 E and G, respectively), whereas for randomly paired-cells, CFSE correlation values dropped between -0.07 for self-renewing cells and 0.18 for differentiating cells (Figure S1 F and H, respectively) indicating no correlation. Those results validated that our general strategy did allow to retrieve accurately generation 1 sister-cells. The same procedure was applied to generation 1 T2EC mother-cells in proliferating phase and in differentiation by sorting the mother-cells either in self-renewing medium or in differentiation-promoting medium.

We further analyzed the T2EC scRNA-seq data quality and reproducibility by characterizing the observed biological process applying UMAP dimensional reduction and projection method (see Methods). As expected, the cells separated based on their differentiation state (Figure S2 A). This observation was validated by a differential expression analysis between the two groups (self-renewing and differentiating cells - Figure S2 B). Genes involved in early erythrocytes maturation, inhibition of differentiation such as *ID2* known to be an erythropoiesis inhibitor in mice [31], *FTH1* and *TMSB4X* known to be expressed in human erythroid progenitors [32] were up-regulated in self-renewing cells while *HBBA, HBAD, HBA1*, genes involved in hemoglobin complex and *TAL1*, erythroid differentiation factor, were up-regulated in differentiating cells, as previously described [28].

### Isolation of second generation cells

Using the T2EC model, we then developed another FACS sorting methodology to retrieve generation 2 sisters and cousin-cells, that is to say the 4 cells resulting from two divisions, both in self-renewing state or in differentiation state. To record cells genealogies, we used different cell-tracers to achieve fluorescent barcoding of cells families and we stained the cells sequentially to retrieve both cousins relationships and sisters relationships within different families (Figure 3).

**Figure 3:**
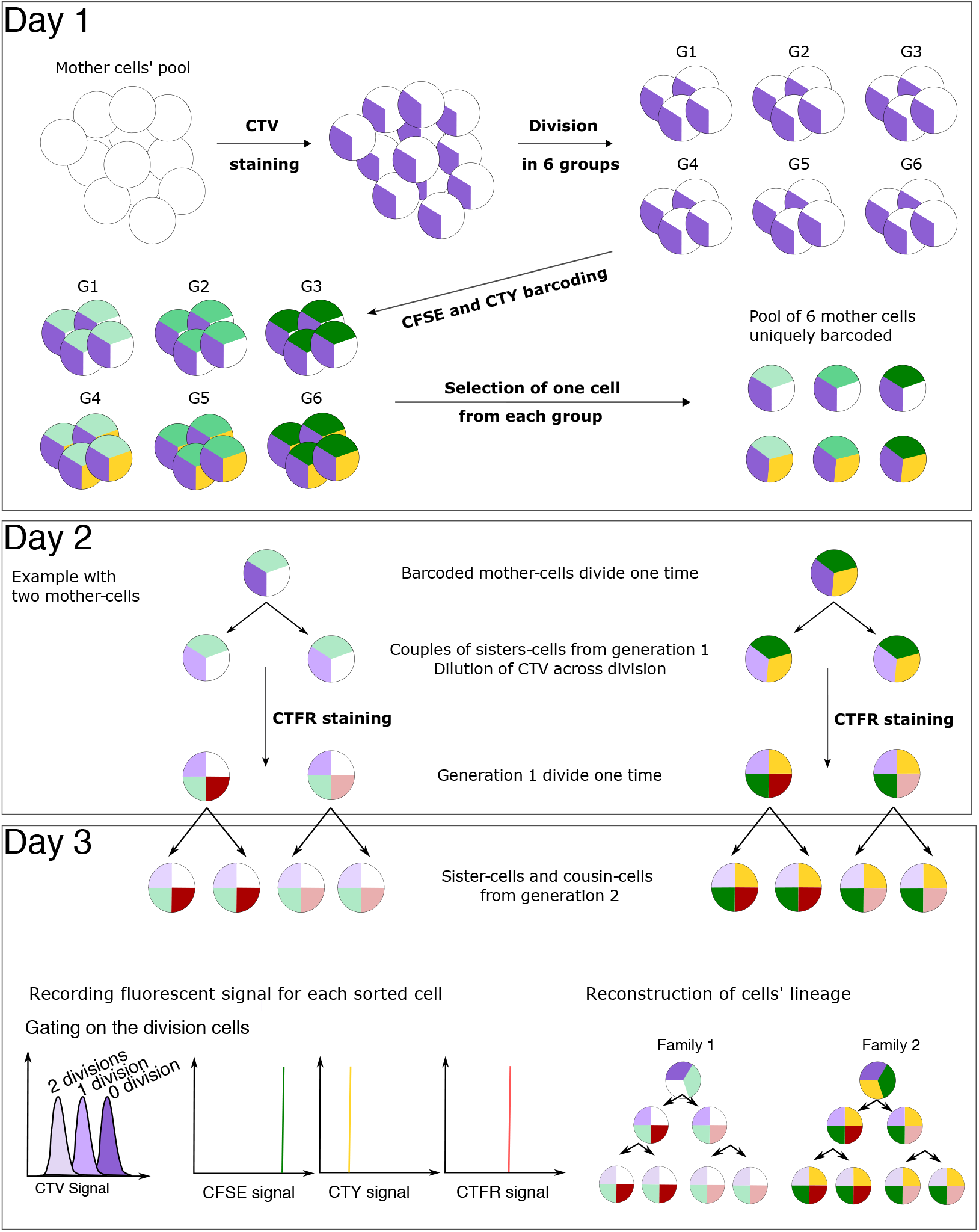
General labelling strategy for generation 2 T2EC cells identification On day 1, a population of mother cells was stained using CTV. The CTV positive population was split into 6 subgroups, each group was barcoded with a unique combination of CFSE and CTY concentration to achieve fluorescent barcoding (6 different barcodes). One mother cell from each group was then recovered and pooled together in a well to be cultured for around 24hrs (6 mother cells with a unique fluorescent barcode). At day 2, after the first division, a fourth dye, CTFR, was added to stain sister-cells with a different intensity in order to be able to discriminate the cells relationship after the next division. On day 3, cells which underwent 2 divisions, determined by the intensity of CTV, were sorted into single-cells and fluorescent intensities were recorded for CTY, CFSE and CTFR signals. Finally, a dedicated script was used to infer the relationships of cells based on the fluorescent intensities (see Methods).

Briefly, a small number of mother-cells was stained such as every mother-cell carried a unique fluorescent barcode. Each fluorescent barcode consist in a combination of CTY and CFSE at different intensities, leading to 6 different barcodes. This barcode is passed along to the mother cells progeny over two cell generations to allow a good discrimination of cells families. One mother cell from each barcode was isolated by FACS in a single well of a culture plate. After the first cell division, another cell-tracer was added to discriminate sister-cells within the cousin groups. After the second cell division, the cells (generation 2) were sorted in lysis buffer containing scRNA-seq primers of known sequence and the relationships between the cells were recovered using a clustering script developed in our team. Details of the methodology are presented in figure 3 and in the Methods section. Further viability analysis was performed and showed that the staining strategy did not compromise cells physiology (Figure S3).

Using first generation methodologies, we successfully collected 86 CD34+ cells, 60 self-renewing T2EC cells and 64 differentiating T2EC cells encompassing respectively 43, 30 and 32 couples of generation 1 sister-cells. With the second-generation original fluorescent barcoding approach, we collected 8 families of generation 2 self-renewing T2EC cells (32 cells) and 5 families of generation 2 differentiating T2EC cells (20 cells).

### Strategy to evaluate transcriptomic similarities between related cells

We used the Manhattan distance as a metric to evaluate transcriptomic similarities between cells. Manhattan distance is a robust geometric distance and is less sensitive to data sparsity, which is inherent to single-cell transcriptomics data [33].

We anticipated how the distance comparisons would result for each of the hypotheses developed in the introduction.

In the case of hypothesis 1, maintenance of memory, there will be no more transcriptional differences between self-renewing than between differentiating sister-cells. This hypothesis would imply that at the first cell generation, differentiating sister-cells would present a similar distance between each other compared to self-renewing sister-cells. And at the second generation, there would be no difference either between differentiating sister-cells compared to self-renewing sister-cells nor between differentiating cousin-cells compared to self-renewing cousin-cells.

In the case of hypothesis 2, gradual erasure of memory, there will be a continuous and gradual increase in the sister-to-sister differences as differentiation proceeds. Meaning, at the first generation, differentiating sister-cells would present a greater distance compared to self-renewing sister-cells. At the second generation, this distance would increase and would be supported by (1) second-generation differentiating sister-cells presenting a greater distance compared to second generation self renewing sister-cells and (2) second generation differentiating cousin-cells presenting a greater distance compared to self renewing cousin-cells.

In the case of hypothesis 3, instantaneous erasure of memory, there will be very strong transcriptional differences between self-renewing and differentiating sister-cells at the beginning of the differentiation process, with no evolution of those differences thereafter. That is, at the first generation, differentiating sister-cells would present an substantial greater distance between each other compared self-renewing sister-cells. At the second generation, differentiating sister-cells cells would display a similar or smaller distance compared to self-renewing sister-cells and differentiating cousin-cells would present a similar or slightly greater distance compared to self renewing cousin-cells.

### Transcriptomic similarities between generation 1 sister-cells after one division

We started by assessing whether or not generation 1 sister-cells displayed more similar global gene expression levels compared to non related cells. Here non related cells correspond to cells which don’t originate from a common mother-cell. The Manhattan distances were computed between the gene expression vectors of each cell. Gene expression vectors for the 43 couples of CD34+ sister-cells were composed of 83 genes after quality control and data filtering (see Methods). Those genes were either selected for their known function in the early differentiation of hematopoietic cells (64% of them) or randomly chosen (36%) to provide an assessment of the overall transcriptional state of the genome. For the 62 couples of T2EC sister-cells gene expression vectors, we retained 1177 genes after data filtering and normalization of scRNA-seq data (see Methods). We performed the analysis by computing the Manhattan distances between generation 1 sisters and randomly selected non related cell pairs from the same pool of cells (Figure 4 A and B).

**Figure 4:**
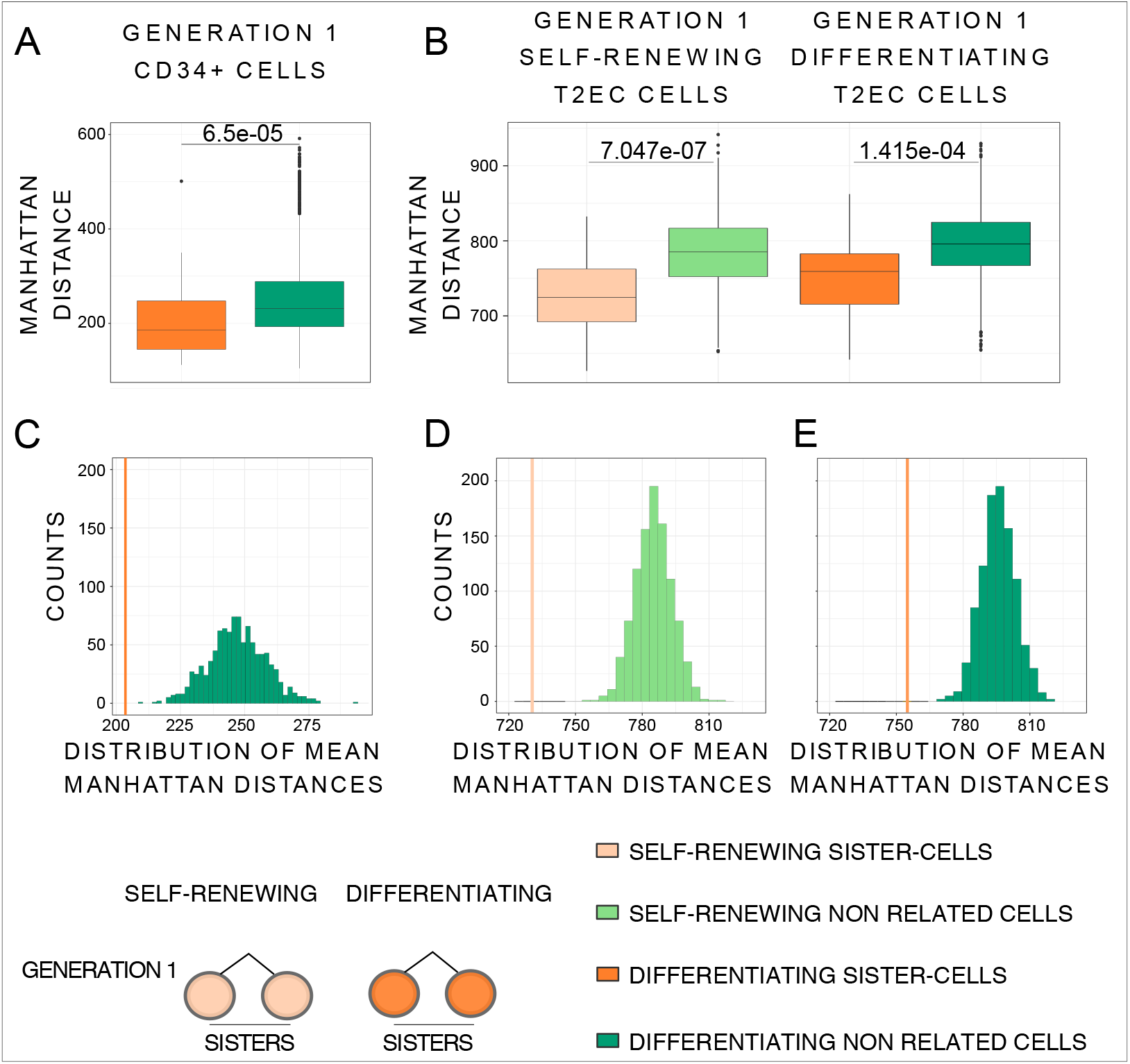
Manhattan distances comparison between generation 1 sister-cells and non related cells. (A) Boxplot of Manhattan distances between the generation 1 CD34+. CD34+ sister-cells (43 couples) are in orange and CD34+ non related cells (3612 couples) in green. Manhattan distances were computed using all the 83 selected genes. Statistical comparison was performed using Wilcoxon test. (B) Boxplot of Manhattan distances between generation 1 T2EC sisters and non related cells. Manhattan distances were computed between all cells from the same biological conditions using all the 1177 selected genes. Self-renewing sister-cells (30 couples) are in light orange and self-renewing non related cells (1740 couples) in light green, differentiating sister-cells (32 couples) are in orange and differentiating non related cells (1984 couples) in green. Statistical comparison was performed using Student t-test. (C) Histograms of mean Manhattan distances of 1000 random draws of distances between 43 CD34+ non related cell pairs (green), compared to the mean distance between the 43 CD34+ generation 1 sister-cells pairs (orange line). (D) Histograms of mean Manhattan distances of 1000 random draws of distances between 30 T2EC self-renewing non related cell pairs (light green histogram), compare to the mean distance between the 30 T2EC self-renewing generation 1 sister-cells pairs (light orange line). (E) Histograms of mean Manhattan distances of 1000 random draws of distances between 32 T2EC differentiating non related cell pairs (Green histogram), compare to the mean distance between the 32 T2EC differentiating generation 1 sister-cells pairs (orange line).

Mean distances were then compared between the two groups (generation 1 sisters and non related cells) for both CD34+ and T2EC cells. For the latter, both self-renewing and differentiating cells were analyzed separately. For both models and in both biological conditions, mean Manhattan distances between generation 1 sister-cells were always significantly smaller than the mean distances between non related cells (Figure 4 A and B - Wilcoxon test for CD34+ cells pvalue = 6.5.10^−5^, Student t-test for self-renewing T2EC cells pvalue = 7.047.10^−7^ and for differentiating T2EC cells pvalue = 1.415.10^−4^).

To ensure that the difference in mean distance observed between generation 1 sisters and non related cells was not an artefact due to difference in sample size, we performed a randomization experiment by bootstrap. Briefly, 43 non related CD34+ cell pairs, 30 non related self-renewing T2EC cell pairs and 32 non related differentiating T2EC cell pairs were randomly drawn from the corresponding groups 1000 times. The mean distance was calculated for each pair and plotted on the histograms shown on figure 4 C, D and E. For both models, and for T2EC in both biological conditions, the mean distance between generation 1 sister-cells was never part of the non related cells mean distances distribution. Those results strongly suggest that the observed difference was genuine and not due to sampling bias.

This is a clear indication that the gene transcription profiles of generation 1 sister-cells in both experimental models are more similar to each-other than to those of non related cells of the same type sharing the same environment and undergoing similar biological processes.

Those results also highlight that differentiating sister-cells from generation 1 display a form of transcriptional memory, which complements previous studies demonstrating a transcriptional memory in self-renewing sister-cells. Focusing on the T2EC model, for which we compared related cells in two cellular states (self-renewing and differentiating), although the difference was borderline non statistically significant (pvalue = 0.06), our results point toward a decrease in transcriptome similarity during differentiation as shown by a higher mean distance value for generation 1 differentiating T2EC sister-cells compared to self-renewing T2EC sister-cells. We wondered whether or not the sister-to-sister cell distance will continue to increase as the differentiation proceeds in the T2EC cells, one generation later.

### Generation 2 cells transcriptomes continue to diverge during differentiation

We generated a second dataset consisting of generation 2 T2EC sisters and cousin-cells (after two cell divisions) using the methodology described above. As scRNA-seq requires the lysis of the cell under investigation, generation 1 data and generation 2 data consist of different cell families and thus cannot be compared to each other so both dataset were treated and analyzed separately (see Methods).

The second generation dataset was composed of 4 cousin-cells per family (8 families of cells in self-renewing and 5 families of cells in differentiation condition), and within the 4 cousins, they consisted of two couples of sister-cells. After data filtering and normalization, we retained 983 genes for subsequent analysis.

Comparison of mean Manhattan distances from those data showed that when comparing conditions, in line with previous results described after one cell generation in figure 4, generation 2 differentiating sister-cells were less close to each other than generation 2 self-renewing sister-cells, although not significantly so (Figure 5).

**Figure 5:**
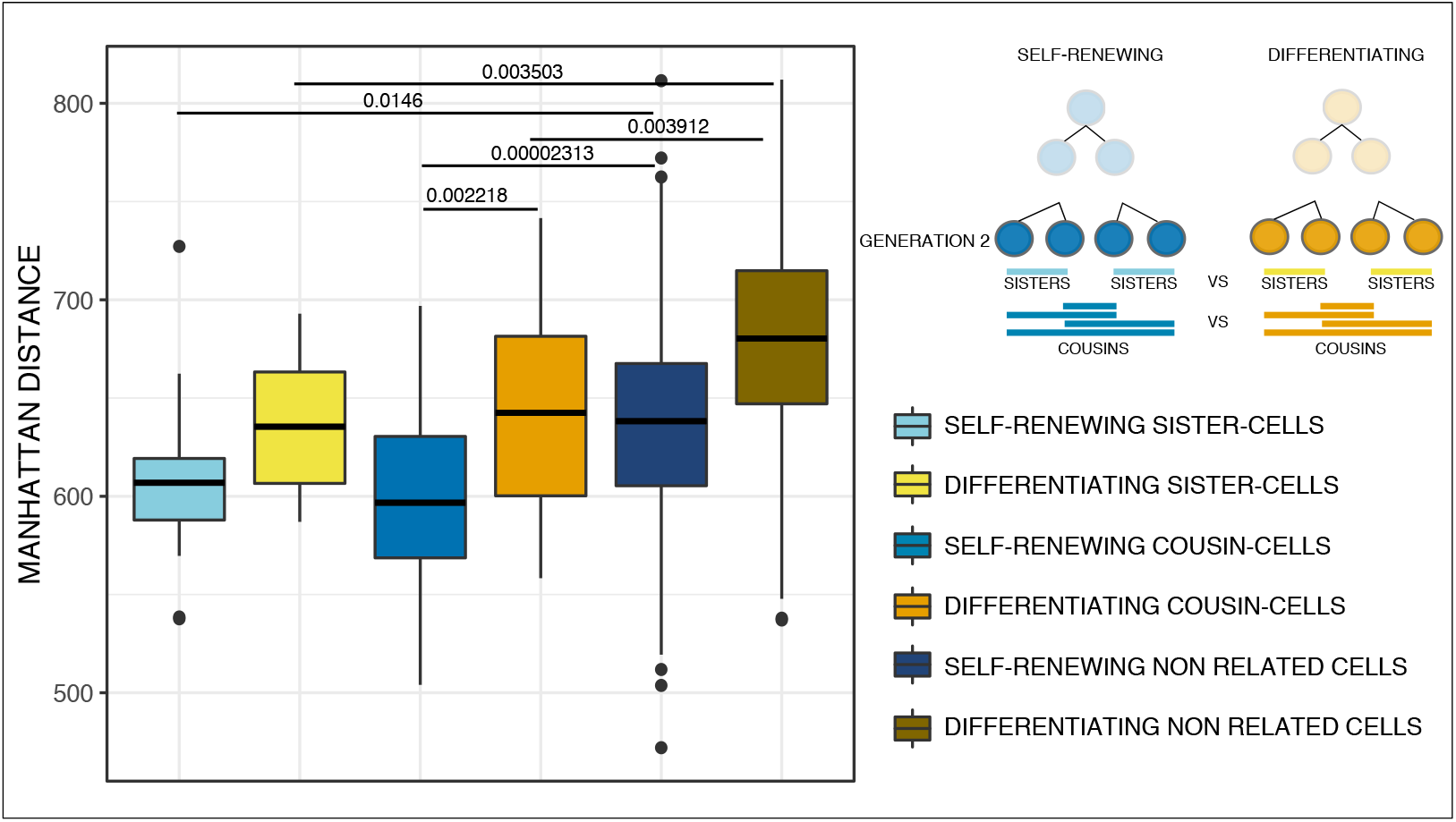
Manhattan distances comparison between generation 2 sisters, cousins and non related T2EC cells. Boxplot of Manhattan distances between generation 2 sisters, cousins and non related T2EC cells. Manhattan distances were computed between all cells (32 self-renewing and 20 differentiating cells) from the same biological condition using the 983 selected genes. Self-renewing generation 2 sister-cells (16 pairs) are presented in light blue, self-renewing generation 2 cousin-cells (32 pairs) are in medium blue and self-renewing non related cells (448 pairs) are in dark blue. Differentiating generation 2 sister-cells (10 pairs) are in yellow, differentiating generation 2 cousin-cells (20 pairs) are in orange and differentiating non related cells (160 pairs) are in brown. Statistical comparisons were performed using Student t-test.

Interestingly, generation 2 differentiating cousin-cells were statistically further apart from the generation 2 self-renewing cousin-cells. Indeed, the average Manhattan distance between generation 2 differentiating cousin-cells was statistically greater than that of generation 2 self-renewing cousin-cells further confirming a decrease in transcriptome similarity during the differentiation process (Student t-test pvalue = 0.002218).

Finally, generation 2 sister-cells, regardless of their biological condition (self-renewing or differentiating for 48hrs), were always closer to each other than randomly paired cells (Figure 5 - Student t-test for self-renewing T2EC cells pvalue = 0.0146 and for differentiating T2EC cells pvalue = 0.003503). Furthermore, the mean Manhattan distances of the generation 2 cousin-cells were also statistically smaller than those of non related cells for both biological conditions, indicating a proximity of transcriptomes which persisted after one more cell generation in both conditions, observed separately (Student t-test for self-renewing T2EC cells pvalue = 0.00002313 and for differentiating T2EC cells pvalue = 0.003912).

### Identification of genes subject to transcriptional memory

We expected that the transcriptomic similarities observed may concern a subset of genes, the “memory genes”, the expression of which would be variable across couples of cells but correlated within couples of sister-cells. Thus, we applied a “gene-wise” approach to identify genes subjected to transcriptional memory using a linear model with random effect and a mixed effects model. For CD34+ cells, memory genes were identified including a sisterhood random effect to capture between-sisters correlation. For T2EC cells, the expression of each gene was modeled by an additive model combining a fixed condition effect (differentiating or not) to account for difference in expression levels and a sisterhood random effect capturing sister-cells correlation. Memory genes were selected by testing for the random effect with a likelihood ratio test comparing the model with and without the sisterhood effect. The test was performed on each gene followed by a Benjamini-Hochberg p-value adjustment for multiple testing [34]. As a negative control, we performed the same test on randomly paired cells, and detected no memory gene (Figure 6).

**Figure 6:**
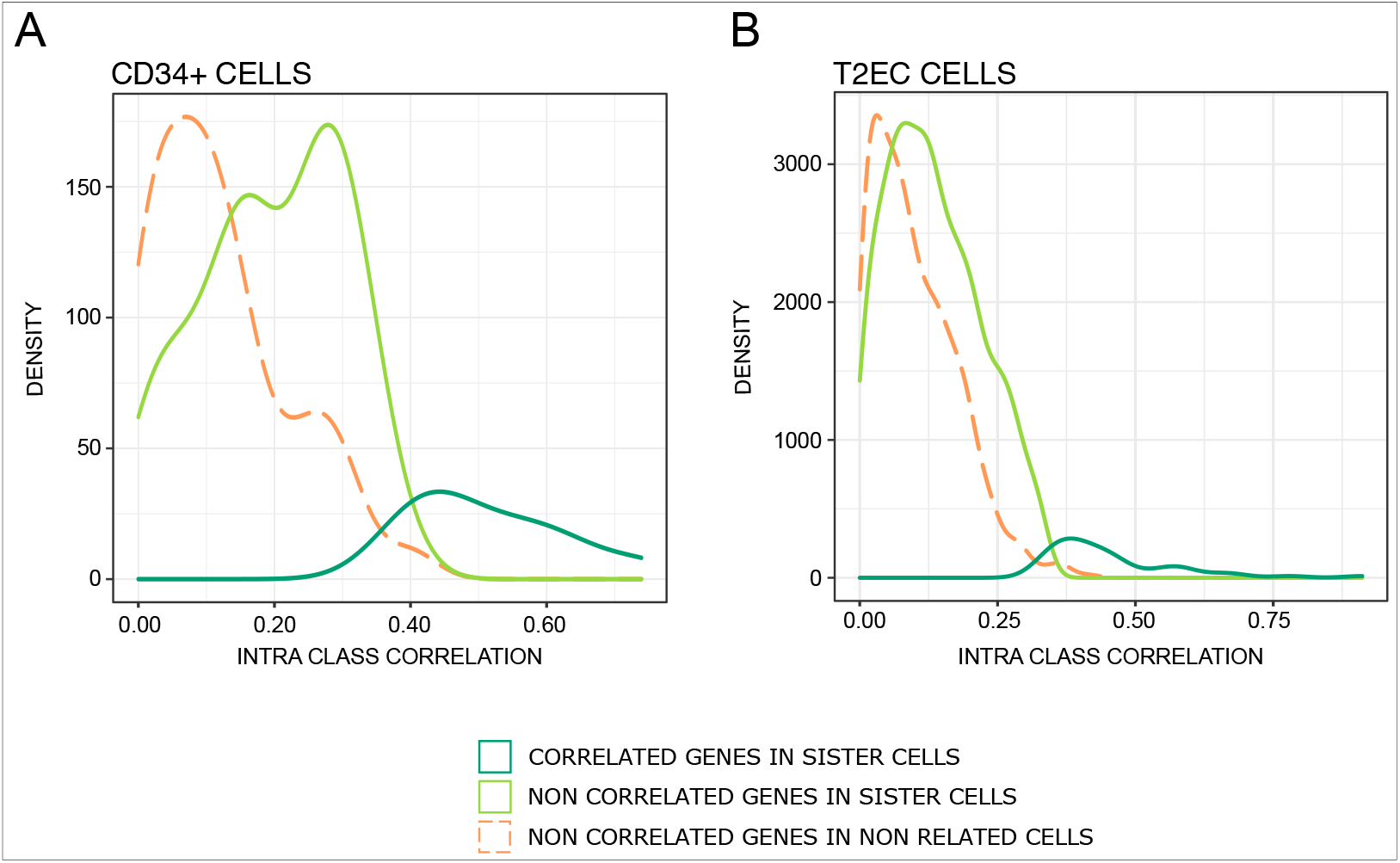
Density plot of genes intra-class correlation in generation 1 sister-cells and randomly paired CD34+ cells (A) and T2EC cells (B). Identification of memory genes using a linear model with random effect (CD34+) and mixed effect model (T2EC). Memory genes are in dark green (11 genes for the 86 CD34+ cells, 55 genes for the 104 T2EC cells, and non significant genes are in light green (72 for CD34+ cells, 1022 for T2EC cells); no memory genes were identified when cells were randomly paired (orange curve).

We detected 10 genes with significant correlation between-sisters in CD34+ cells and 55 genes in T2EC cells (cf. Supplement Table S1 for CD34+ and for T2EC). In CD34+ cells, memory genes were involved in diverse functions, including stemness (*GATA1,CD38, CD133*), differentiation and proliferation (*CD74, ERG, KIT*), metabolism (*BCAT1, HK1*), cytoskeleton (*ACTB*) and tRNA splicing (*C22orf28*). In T2EC, memory genes were involved in erythropoietic differentiation (*HBBA, HBA1, HBAD*, which are hemoglobin subunits, or *RHAG* membrane channel component involved in carbon dioxide transport), chromosome structure (*SMC2, H2AFZ*), ribosomes and translation (*RPS13, RPL22L1, UBA52, EEF1A1*) and metabolism (*GAPDH, LDHA*). One should note that *LDHA* was previoulsy found to also be involved in the erythroid differentiation process [25].

We computed again and compared the Manhattan distances for the T2EC cells between sisters and non related cells using as a vector only the 55 memory genes (Figure 7). As a result, the difference in within-distance between sister-cells and non related cells, in both biological conditions (self-renewing and differentiating), was even more pronounce than when computing the Manhattan distances using all 1177 genes of the scRNA-seq dataset (see above), further confirming that the identified genes are the ones imprinted by the transcriptional memory.

**Figure 7:**
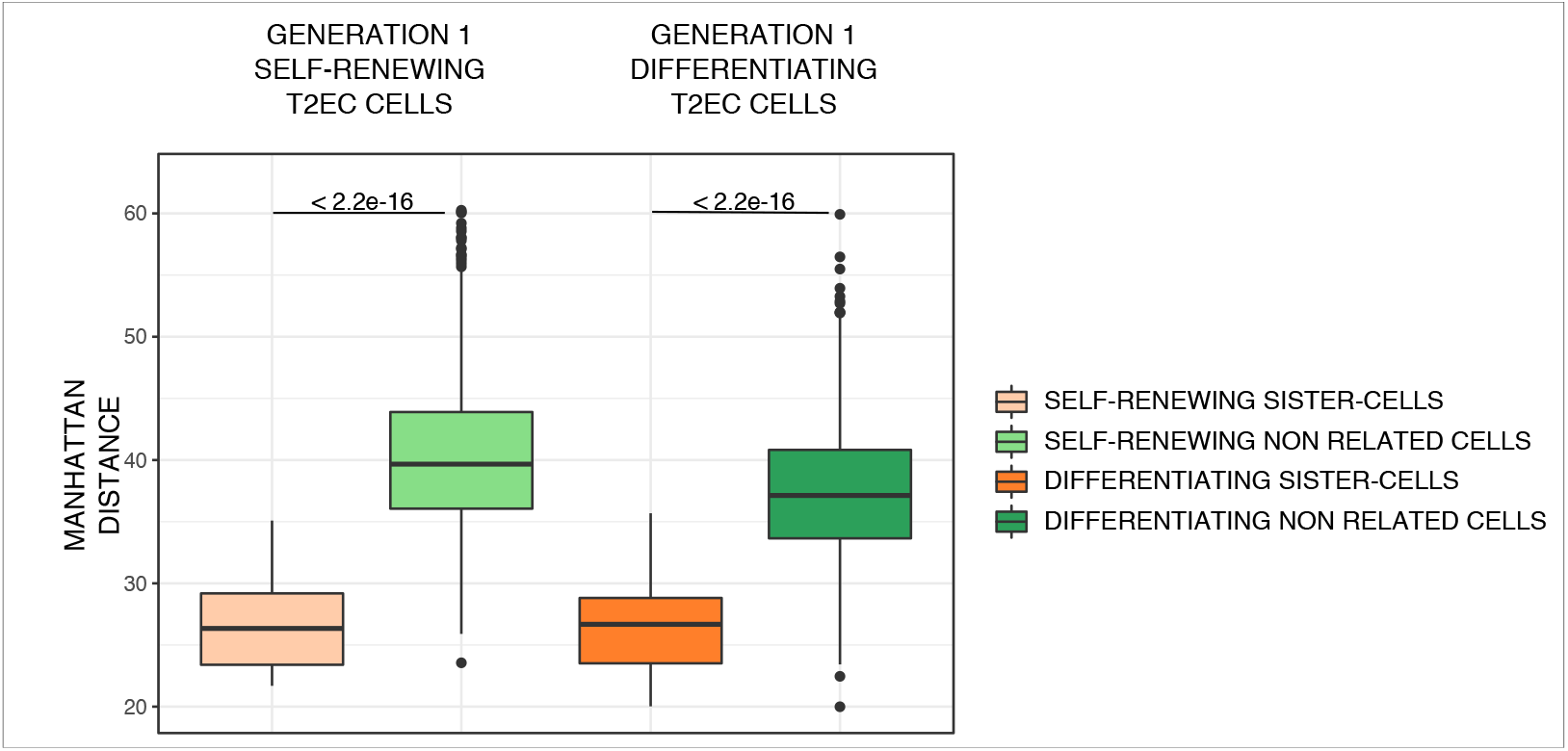
Manhattan distances comparison between generation 1 sisters and non related T2EC cells using the memory genes. The Manhattan distances were computed between all cells from the same biological conditions using all the 55 memory genes. Self-renewing sister-cells (30 couples) are in light orange and self-renewing non related cells (1740 couples) in light green, differentiating sister-cells (32 couples) are in orange and differentiating non related cells (1984 couples) in green. Statistical comparison was performed using Wilcoxon test.

To validate our findings, we also checked if these memory genes were not only genes associated high mRNA half-life. We crossed our gene list to a previously published dataset which evaluated half-life duration of genes during T2EC differentiation using RT-qPCR [35]. We were able to compare the half-life duration of 6 memory genes and found that 4 of them have a relatively long half-life but 2 of them have a quite short half-life (Figure 8A). Furthermore, other genes with longer half-life were not identified by the model as memory genes. Thus, half-life duration could not be the only cause of memory.

**Figure 8:**
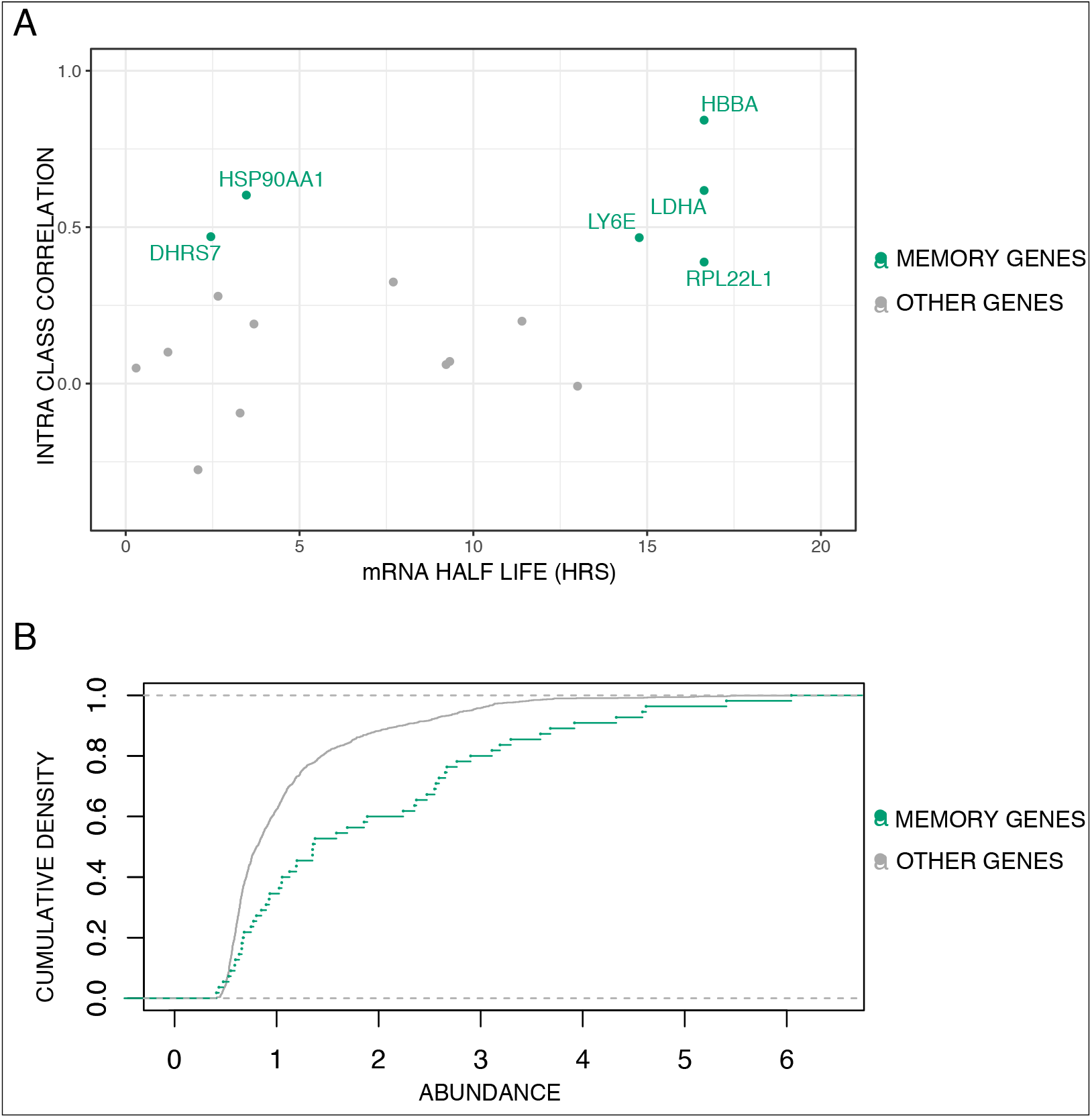
T2EC Memory genes characteristics. (A) mRNA half-life of memory genes and other genes present in the scRNA-seq dataset evaluated at 24hrs post differentiation induction ([35]) vs their Intra Class Correlation value extracted from the mixed effects model. (B) Cumulative empirical distribution graph of transcripts abundance of the 55 Memory genes in the dataset compared to total genes (1177) of scRNA-seq data.

We also questioned the relationship between the level of expression of a gene and its belonging to the memory genes class. 1000 bootstrap distribution analysis of the abundance of the 55 memory genes compared to the abundance of 55 randomly drawn genes showed an enrichment for higher abundance of the 55 memory genes (Figure 8B - kolmogorov-Smirnov test pvalue = 0.01672). We therefore can not exclude that part of the memory is due to high level expression for at least some memory genes and could be related to synthesis and degradation dynamics. However, this result was expected because to prevent false correlation that would be due to high numbers of zeros in expression value of lowly expressed genes between sister-cells, we selected genes with mid to high-level of expression in our scRNA-seq data set (see Methods). Finally, we didn’t regress cell-cycle effects on our data, due to the fact that cell-cycle is not as well described in chicken cells as it is in mammalian cells, and thus cannot exclude that the sister-to-sister resemblance may, in part, be a consequence of the sister-cells being at similar state in the cell-cycle. However, while we found a GO term “cell-cycle” enrichment in the 1177 selected genes, no cell-cycle related genes were identified as memory genes, leading us to believe that cell-cycle is not the main driver of this transcriptional memory.

## Discussion

In the present study, we questioned the interplay between the transcriptional memory and the gene expression variability which characterizes differentiation processes.

We developed two experimental frameworks to recover sister-cells (Generation 1) and one experimental framework to recover cousin-cells (Generation 2) transcriptomes while preserving the information about their lineage at the resolution of the cell division. We analyzed the transcriptomes of related cells from two different cell differentiation systems using two different single-cell transcriptomics technologies.

Comparison of global transcriptomic state, using Manhattan distances, showed that differentiating generation 1 sister-cells (both CD34+ cells and T2EC cells) transcriptomes are globally significantly more similar between each other than between non related cells.

In our controlled differentiation model (T2EC cells), we observed after one cell division (generation 1), a greater mean distance for differentiating sister-cells compared to self-renewing sister-cells. Moreover, the difference becomes significant after a second division (generation 2), showed by differentiating cousin-cells presenting a significantly higher distance than self-renewing cousin-cells. Those results showed that during cell differentiation, related cells deviates faster from each other than during self-renewing divisions.

Mixed models further highlighted that some genes have their expression statistically correlated between sister-cells while none were found between non related cells. We qualified those genes as “memory genes” and obtained evidence that they weight out the transcriptomic resemblance observed between sisters and cousin-cells. However, the mechanisms leading to a more correlated expression between related cells for those genes remain to be investigated.

In the introduction, we formulated 3 hypothesis on the possible evolution of the transcriptional memory upon differentiation induction (Figure 1). Our results therefore support the second hypothesis: upon differentiation induction, transcriptional memory is continuously and gradually erased eventually reconstituting, at the clonal scale, the variability observed in the initial population.

While our experimental methods allow to preserve genealogical cell information for two generations, everything happening later is presently out of reach. We therefore are currently developing a microfluidics-based approach, consisting in a microfluidics chip coupled to scRNA-seq, which could be used on non-adherent cells to investigate cellular memory for several (more than 2) generations. Recently, a study based on a complex cell-tracking system combining time-lapse microscopy, antibody-based cell isolation and scRNA-seq on robotically-isolated cells has been used to address the question of asymetric division [36]. While the question is different from ours, the approach could be considered to investigate longer genealogies but it would require complex equipments and antibodies against chicken cells in order to track division.

In order to explain the existence of memory genes as we (this work) and others [5–8] have described, one need to assume that a significant fraction of those mechanisms must “survive” the mitosis, i.e. be transmitted through the dramatic epigenomic and cellular rearrangements involved in the cell division process. If one assumes that the GRN state is essentially characterized by protein quantities, then it is easy to see that it will be pass through, at least for the proteins with a sufficiently long half life [15]. Reestablishment of the epigenetic marks [37] and of genomic structure [38] after a division process have also been documented.

It has recently been described that the persistence of a low level of transcription throughout the mitosis might at least partly explain how transcriptional memory can be maintained. It would be interesting in that regard, to assess the overlap between our memory genes and these genes for which the mitotic transcription can be detected using UEseq in mitotic chromosomes [39].

Differentiating division is a specific challenge since at each division a subtle combination of changes and stability must be imposed. In this respect one can see the bookmarking process [40] as a stabilizing process, whereas the increase in gene expression variability [11–19] will affect the GRN state and therefore will tend to modify gene expression burst parameters. In fact, at the single cell level, gene expression is in essence a probabilistic process that is characterized by a given burst frequency and burst size [41]. The mechanisms regulating this bursting process are still a matter of debate [42, 43], but are usually thought to involve: 1) the state of the underlying Gene Regulatory Network (GRN) [44]; 2) the state of the chromatin, a.k.a. the epigenetic marks [7, 8], and 3) the genomic 3D state [45]. Of course none of these mechanisms operate in isolation and more integrated mechanisms, like the metabolism, are also key players in the burst properties of transcription (see e.g. [46]).

It is interesting to note that our two model systems do behave quite differently in regard to the division process. The initial stages of T2EC erythrocytic differentiation have been shown to result in an increase of the proliferation rate due to a shortening of the G1 period [23]. This is in sharp contrast with the observation that the CD34+ first division occurs after an unusually long cell cycle that lasts on average more than 55 hrs [15]. It could therefore be that the molecular mechanisms linking cell division and differentiation might be quite different in the two cell types, although the final result will be similar: cellular memory will show a high level of robustness in front of the cellular state change associated with the differentiation process.

Finally, it is tempting to speculate that the observed burst in entropy at the beginning of the differentiation sequence is helping the differentiating cells to overcome a memory process that is meant to prevent changes in cellular identity.

## Material and methods

### Cell culture

Human hematopoietic CD34+ cells were purified from umbilical cord blood from three anonymous healthy donors. First, mononuclear cells were isolated by density centrifugation using Ficoll (Biocoll, Merck Millipore). CD34+ cells were then enriched by immunomagnetic beads using the AutoMACSpro (Miltenyi Biotec). Cells were frozen in 90% fetal bovine serum (Eurobio) 10% dimethylsulfoxide (Sigma) and stored in liquid nitrogen. After thawing, cells were grown in prestimulation medium made of Xvivo (Lonza) supplemented with penicillin/streptomycin (respectively 100U/mL and 100µg/mL - Gibco, Thermo Scientific), 50 ng/ml h-FLT3-ligand, 25 ng/ml h-SCF, 25 ng/ml h-TPO, 10 ng/ml h-IL3 (Miltenyi) final concentration as previously described [15]. Cells were cultured in a 96-well plate at 185 000 cells/mL during 24hrs in a humidified 5% CO2 incubator at 37°C before proceeding to mother cells isolation.

Cell population mortality was assessed by counting dead and living cells from the different time points and conditions after Trypan blue staining and using a Malassez cell.

T2EC cells were extracted from 19-days-old SPAFAS white leghorn chicken’s embryos’ bone marrow (INRA, Tours, France). Cells were grown in LM1 medium (*α*-MEM, 10% Fetal bovine serum (FBS), 1 mM HEPES, 100 nM *β*-mercaptoethanol, 100 U/ mL penicillin and streptomycin, 5 ng/mL TGF-*α*, 1 ng/mL TGF-*β* and 1 mM dexamethasone) as previously described [23]. T2EC cells differentiation was induced by removing LM1 medium and placing the cells into DM17 medium (*α*-MEM, 10% fetal bovine serum (FBS), 1 mM Hepes, 100 nM *β*-mercaptoethanol, 100 U/mL penicillin and streptomycin, 10 ng/mL insulin and 5% anemic chicken serum [24]).

### Manual strategy for CD34+ sister-cells isolation

Mother cells were isolated using a SmartAliquotor (iBioChips). It consists of a polydimethylsiloxane chip divided into 100 wells (2*µ*L per well, 1.8mm of diameter) connected by microchannels to an insertion hole in the center. This system allows to physically isolate cells while sharing the same medium. 200*µ*L of cell suspension at 1000 cells/mL were injected in the chip through the injection plug and cells were randomly divided into the wells. Air bubbles were removed with sterile tips. Using a standard confocal microscope, wells containing lonely cells were listed. 20mL of prestimulation medium (see Cell culture part for composition) were added to avoid evaporation and cells were incubated at 37°C in a humidified 5% atmosphere during 24 to 48hrs. Listed wells were regularly checked with standard confocal microscope to identify cell division. Sister-cells were manually collected under biological safety cabinet to keep sterile conditions and avoid impurities to fall in the culture dish. A micromanipulator connected to a flexible microfluidic capillary filled with PBS and ending in a 2*µ*L glass microcapillary was used. Individual collected cells were immediately inserted into 5*µ*L of lysis buffer (Triton 4% (Sigma), RNaseOUT Recombinant Ribonuclease Inhibitor 0.4U/µL (Thermo Scientific), Nuclease free water (Thermo Scientific), Spikes 1 and 4 (Fluidigm C1 Standard RNA Assays)) and kept on dry ice to preserve RNA. Particular attention has been given to preserve cells integrity. Samples were kept at -20°C until further sc-RT-qPCR analysis.

### FACS-oriented strategy for T2EC sister-cells isolation

Mother cells were stained using CFSE (Cell Trace CFSE Cell Proliferation kit Thermofisher), 5×10^5^ cells were placed in a 60mm plate in 5mL of culture medium mixed with 5*µ*L of CFSE at 5 mM (final concentration 5*µ*M) and incubated at 37°C for 30min. Cells were then centrifuged at 20°C, 1500rpm for 5min. Medium was discarded and cells were resuspended in 5mL fresh medium. CFSE stained mother cells were then isolated using the CellenONE X1 (CELLENION) at CELLENION core facility (Lyon, France). A gating based only on morphological criteria (diameter, elongation and circularity) was performed to select single living cells. Selected single cells were sorted in a 384-well plate containing 10*µ*L of culture medium (either self-renewing medium LM1 or differentiation-inducing medium DM17). The plate was then kept in an incubator under 5% CO2, 37°C for at least 20hrs to allow one cell division. Each well of the 384-well plate was manually checked under a regular inverted microscope to identify cells that had undergone one cell division (presence of cell doublets). Each doublet was then harvested and placed in a FACS polypropylene tube containing 80*µ*L of warm culture medium. Tubes containing cell doublets were kept at room temperature throughout the sorting process and were briefly vortex immediately before loading into the sorter. Prior settings consisted in analysing the CFSE positive population, the CFSE negative population and the culture medium. No fluorescent signal was ever detected in medium or in negative population (Figure S1 A-B self-renewing medium and C-D differentiation medium) indicating that only cells of interest ever gave CFSE positive signal. Cells were sorted at 20 PSI through a 100 *µ*m nozzle on an FACS AriaII (BD). Gating was performed on FSC-A/SSC-A to capture live cells, SSC-H /SSC-A to capture single cells, and CFSE positive cells with yield, purity and phase mask of 32, 0, 0 respectively. Those parameters were chosen because cell density being very low (2 cells per tube), the probability of the two cells being in two consecutive drops was extremely low. Furthermore, those parameters are very conservatives and thus probability of the cell not being sorted is also very low. Cells were isolated in 4*µ*L of lysis buffer in PCR tubes containing cell barcode primers. Tubes were frozen in dry ice directly after sorting to prevent any degradation of the samples.

### FACS-oriented strategy for T2EC cousin-cells isolation

#### Fluorescent barcoding for lineage tracing

On the first day, 1×10^6^ mother cells were labelled with 0.5*µ*M CTV (Cell Trace Violet Cell Proliferation kit Thermofisher) for 20min at 37°C in PBS, then 5mL of medium was added for 5min to dilute the fluorescent molecules. The cells were centrifuged for 5min at 1500rpm at 20°C, resuspended and then separated into 6 tubes (2×10^5^ cells per tube) and resuspended in 1mL per tube. Each sample was labelled with a different concentration of CFSE (3-point range of 5*µ*M, 2.187*µ*M and 0.312*µ*M) plus or minus CTY (10*µ*M - Cell Trace Yellow Cell Proliferation kit Thermofisher) for 30min at 37°C in PBS. Each condition was centrifuged for 5min at 1500rpm at 20°C and resuspended in 1mL of fresh medium. The different concentrations and combinations were optimised so that even after two cell divisions, the barcodes will be different enough to differentiate the cell clones. Cells were plated in a 6-well plate and kept in culture conditions until sorting (in an incubator 37°C, 5% CO2). Cells were were stored at 37°C throughout the sorting process and sorted at 20 PSI through a 100 *µ*m nozzle on an FACS AriaII (BD). The sorting strategy was done using single-labelled cell populations (CFSE, CTY, CTV and negative), then gating was performed on FSC-A/SSC-A to capture live cells, SSC-H /SSC-A to capture single cells, and CTV positive cells. One cell from each subgroup (6 cells total) was isolated in a well of a 96-well plate which contained 500 non-labelled feeder cells in either self-renewing medium or differentiating medium through a 100*µ*m nozzle with yield, purity and phase mask of 0, 32, 16 respectively (single-cell mask). A well then contained 6 mother cells, each one labelled with a unique fluorescent barcode and the feeder cells. The plate was then put back in culture conditions (in an incubator 37°C, 5% CO2).

CTFR (Cell Trace Far Red Proliferation kit Thermofisher) labelling was performed 20hrs after mother cells sorting, in the plate, so that the cells had time to divide once. The staining was made as heterogeneous as possible, thanks to the feeder cells but also by using very low concentrations of dye and for a very short amount of time. Indeed, 0.37*µ*M of CTFR (Cell Trace Far Red Cell Proliferation kit Thermofisher) was added to each sample (in approximately 50*µ*L of medium), and then 100*µ*L of medium was added to dilute the dye. The plate was centrifuged for 5min at 200G, then 120*µ*L of medium was removed and 50*µ*L of new medium added to each labelled well. This heterogeneous CTFR staining will allow to discriminate the next division meaning within the 4 cousin-cells, how they are paired two by two. Indeed, each daughter-cell will receive a unique intensity of CTFR dye which will be discriminating after one more cell division. Cells were kept in culture conditions for an additional 20hrs (in an incubator 37°C, 5% CO2).

On the third day, after the second division, the content of the wells containing the cousin-cells were transferred into polypropylene FACS tubes and briefly vortexed immediately before loading into the sorter. The sorting strategy was done using single-labelled cell populations (CFSE, CTY, CTV, CTFR and negative), then gating was performed on FSC-A/SSC-A to capture live cells, SSC-H /SSC-A to capture single cells, and CTV positive corresponding to the second division peak and exclude feeder cells. Cells were sorted on a FACS AriaII (BD) at 20 PSI through a 100*µ*m nozzle with yield, purity and phase mask of 32, 16, 0 respectively, in PCR tubes containing lysis buffer (0.2% Triton (Sigma Aldrich), 0.4 U/*µ*L RNaseOUT (Thermofisher Scientific), 400nM RT primers (Sigma Aldrich)) and scRNA-seq primers. The fluorescent intensities for CFSE, CTY and CTFR were recorded for each cell to further reconstruct relationships between the cells using our clustering algorithm.

#### Cousin-cells identification

Clustering was performed using the R mclust package [47] (version 5.4.10 - https://gitbio.ens-lyon.fr/LBMC/sbdm/sister-cells commit 76615c6e). This clustering script finds the genealogical relationships between cells in two steps. First, cousin-cells are grouped together by their fluorescent barcode, determined by the CTFE and CTY fluorescent intensity values. Thus, if two, three or four cells have the same CFSE and CTY intensities levels they will be considered as cousins. In a second step, we select the groups for which the 4 cousin-cells were sorted in the plate, then the program identifies the two pairs of sisters within the 4 cousins. To do this, the median CTFR intensity is calculated, then the two cells that have intensity values higher than the median are matched, and the other two that have lower intensity values are matched together. Finally, when sorting, we used an index sorting option, which allows us to know in which well of the plate each cell was sorted. With this position information, our analysis program returns the position of the retained cells, i.e. the cells belonging to the cousin groups for which the 4 cells were successfully isolated in the lysis plate.

### sc-RT-qPCR data generation

#### sc-RT-qPCR one step

Lysed cells were heated at 65°C during 3 minutes for hybridization with RT primer and immediately transferred into ice. 7*µ*L of RT-PCR mix (Superscript III RT/platinium Taq 0,1*µ*L (Invitrogen), Reverse and Forward primers and spikes at 1,33*µ*M final concentration and homemade 2X reaction Mix (120mM Tris SO4 pH=9, 2.4 mM MGSO4, 36mM (NH4)2SO4, 0.4mM dNTP)) were added to each well before launching of reverse transcription and PCR run on thermocycler (Program : 50°C 15min - 95°C 2min - 20 cycles 95°C 15sec/60°C 4min - Hold 4°C). 3*µ*L of exonuclease mix (Exonuclease I 1.6U/mL (NEB), Exonuclease buffer 1X (NEB), Nuclease free water (Thermo Scientific)) were added and samples were incubated for a digestion run on thermocycler (Program : 37°C 30min - 80°C 10min). Pre-amplified samples were diluted five times in TE low EDTA (10mM Tris, 0.1mM EDTA, pH=8) and kept at -20°C for one night before qPCR.

#### qPCR with Fluidigm Biomark technology

3,15*µ*L of pre-amplified samples were distributed into a 96-well plate and 3,85*µ*L of qPCR mix (Sso EvaGreen Supermix with Low ROX (Bio-Rad)+ 20X DNA binding dye sample loading reagent) were added to each well. Simultaneously, a 96-well plate with primer mix (forward and reverse primers and spike at 2*µ*M final concentration, 2X Assay Loading readent, TE low EDTA) was prepared. The microfluidigm chip was primed with injection oil using the IFC Controller HX (Fluidigm). 5*µ*L of primers and 5*µ*L of samples were loaded in the dedicated wells of the chip. Air bubbles were removed with a needle. Samples and primers were mixed in the IFC Controller HX (Fluidigm) with the loading program. The chip was then transferred in the Biomark HD system (Fluidigm) for qPCR with “HE 96×96 PCR+Melt v2.pcl” thermal cycling protocol with auto exposure.

#### Quality control and Normalization

Ct values obtained from the Biomark HD System (Fluidigm) were exported as excel files and quality control was manually done. For each gene, “failed” quality control readings identified by the Fluidigm software were removed. Four negative controls (mix of water and lysis buffer) were used to detect unwanted amplification and the associated genes were also removed. Finally, two externally added controls (spike 1 and spike 4, Fluidigm) were used to control amplification consistency. Filtered data frame was then imported into R (version 4.2.0) for normalization to remove amplification bias (https://gitbio.ens-lyon.fr/LBMC/sbdm/sister-cells commit 45a65972). For each cell, expression values were calculated by subtracting the gene Ct value from the geometric mean of Ct values from spike 1 and spike 4 of the corresponding well. Then, an arbitrary differential cycle threshold value of -22 for null signal (corresponding to a Ct value of 30) was assigned for all genes with a Ct value less than -22.

### scRNA-seq data generation

#### scRNA-seq libraries preparation

Subsequently to sister or cousin-cells isolation, we performed single cell RNA sequencing (scRNA-seq) using a modified version of the Mars-seq protocol [26] published here [28]. This specific protocol of scRNA-seq allowed us to know in advance which cell barcode would be carried by each cell and thus preserving the genealogy information of the cells. Briefly, Reverse Transcription (RT) was performed so every mRNA of the cells were tagged with a combination of unique cell barcode and a 8pb random UMIs sequence for further demultiplexing. After barcoding, all mRNA were pooled and second DNA strand were synthetized. Amplification was done over night using In Vitro Transcription (IVT) to obtain a more linear amplification. A second barcode was added by RT to identify plates. Libraries were amplified by PCR and Illumina primers were added.

#### Sequencing

Libraries were sequenced on a Next500 sequencer (Illumina) with a custom paired-end protocol (130pb on read1 and 20pb on read2) and a depth of 200 000 raw reads per cell.

#### Data preprocessing

Fastq files were pre-processed through a bio-informatics pipeline developed in our team on the Nextflow platform [48], available here https://gitbio.enslyon.fr/LBMC/sbdm/mars_seq and published here [28]. Briefly, the first step removed Illumina adaptors. The second step de-multiplexed the sequences according to their plate barcodes. Then, all sequences containing at least 4T following the cell barcode sequence and UMI sequence were kept. Using UMItools whitelist, the cell barcodes and UMI sequences were extracted from the reads. The cDNA sequences were then mapped on the reference transcriptome (Gallus GallusGRCG6A.95 from Ensembl) and UMIs were counted. Finally, a count matrix was generated for each plate.

#### Quality control and data filtering

All analysis were carried out using R software (version 4.1.2; [49]) and are available on the following git repository https://gitbio.ens-lyon.fr/LBMC/sbdm/sister-cells. For the sister-cells dataset, cells were filtered based on several criteria: reads number, genes number, counts number and ERCC content. For each criteria the cut off values were determined based on SCONE [50] pipeline and were calculated as follows:

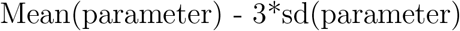

We then removed orphan cells, meaning cells which sister was not present in the dataset. After filtering, we kept 60 undifferentiated cells (30 couples) and 64 differentiating cells (32 couples). For the cousin-cells dataset we performed the same filtering strategy and kept only cell groups which contained the 4 cousin-cells. After filtering we kept 32 undifferentiated cells (8 groups of cousins) and 20 differentiating cells (5 groups of cousins). Based on [51] work, genes were kept in the data set if in mean present in every cell. After applying this filter, we kept 1177 and 983 genes for the sister-cells dataset and the cousin-cells dataset respectively.

#### Normalization

Filtered matrix were normalized using SCTransform from Seurat package (version 1.6 [52] - https://gitbio.ens-lyon.fr/LBMC/sbdm/sister-cells commit 945aaca7 and 94f13467) and were corrected for batch effect, day of isolation effect, medium effect and sequencing depth effect. Both datasets (sister-cells and cousin-cells) were processed independently.

## Bioinformatics analysis on R

All analysis were carried out using R software [49] (version 4.1.2 for T2EC and version 4.2.0 for CD34+). Plots were performed ggplot2 package (version 3.3.6).

### Dimensional reduction

UMAP dimension-reduction and visualization were performed using UMAP package (version 0.2.8.0; [53]).

### Manhattan distance computation

Distances were computed on normalized matrix between all cells using dist function from R software. Distances between sister-cells were extracted and compare to the same number of randomly chosen distances of non related cells. 1000 bootstraps were performed this way. Mean comparison was performed using Student t-test or Wilcoxon test when Student t-test was not applicable (https://gitbio.ens-lyon.fr/LBMC/sbdm/sister-cells commit 8417545d and 45a65972).

### Linear model with random variable and Mixed effects model

Linear model with random variable and Mixed effects model analysis were performed using lme4 R package (version 1.1-29 - https://gitbio.ens-lyon.fr/LBMC/sbdm/sister-cells commit c24fa472). The models were defined as followed:

Mixed effect Model definition :

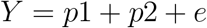

Linear Model with random variable definition :

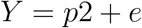

where Y is the mean expression of each gene, p1 is the fixed effect and p2 is the random effect. Here, p1 corresponds to the biological condition and can take two values (undifferentiated and differentiating) and p2 is the sorority effect. Two sister-cells have the same discrete value. And e is the error of the model. Null models are the above model but without the random effect e.g. the sorority effect. Genes were selected based on a significant adjusted BH p-value after performing a likelihood ratio test between the model and the null model.

## Acknowledgements

We gratefully thank all members of SBDM team for very fruitfull discussions, suggestions and commentaries on our project. We thank the computational center of IN2P3 (Villeurbanne/France) and Pôle Scientifique de Modélisation Numérique (PSMN, Ecole Normale Supérieure de Lyon) where computations were performed. We thank the BioSyL Federation and the LabEx Ecofect (ANR-11-LABX-0048) of the University of Lyon for inspiring scientific events. We acknowledge the contributions of the CELPHEDIA Infrastructure (http://www.celphedia.eu/), especially the center AniRA in Lyon.

## Declarations

### Ethics approval and consent to participate

Human cord blood (UCB) was collected from placentas and/or umbilical cords obtained from AP-HP, Hôpital Saint-Louis, Unité de Thérapie Cellulaire, CRB-Banque de Sang de Cordon, Paris, France (Authorization number: AC-2016-2759) or from Centre Hospitalier Sud Francilien, Evry, France in accordance with international ethical principles and French national law (bioethics lawnn°2011-814) under declaration N° DC-201-1655 to the French Ministry of Research and Higher Studies.

### Consent for publication

Not applicable

### Availability of data and materials

Data tables are supplied as supplements files. scRNA-seq data are available in SRA repository under the BioProject accession PRJNA882056 and BioSample accessions SAMN30926136 and SAMN30926137. Embargo will be released upon publication.

R codes are available at : https://gitbio.ens-lyon.fr/LBMC/sbdm/sister-cells Embargo will be released upon publication.

## References

1. Miura, H. & Hiratani, I. Cell cycle dynamics and developmental dynamics of the 3D genome: toward linking the two timescales. Curr Opin Genet Dev 73, 101898. issn: 1879-0380 0959-437X (2022).

2. Sigal, A., Milo, R., Cohen, A., Geva-Zatorsky, N., Klein, Y., Liron, Y., Rosenfeld, N., Danon, T., Perzov, N. & Alon, U. Variability and memory of protein levels in human cells. Nature 444, 643–646. issn: 0028-0836, 1476-4687 (Nov. 2006).

3. Schwanhäusser, B., Wolf, J., Selbach, M. & Busse, D. Synthesis and degradation jointly determine the responsiveness of the cellular proteome: Insights & Perspectives. BioEssays 35, 597–601. issn: 02659247 (July 2013).

4. Corre, G., Stockholm, D., Arnaud, O., Kaneko, G., Viñuuelas, J., Yamagata, Y., Neildez-Nguyen, T. M. A., Kupiec, J.-J., Beslon, G., Gandrillon, O. & Paldi, A. Stochastic Fluctuations and Distributed Control of Gene Expression Impact Cellular Memory. PLoS ONE 9 (ed MacArthur, B. D.) e115574. issn: 1932-6203 (Dec. 22, 2014).

5. Kimmerling, R. J., Lee Szeto, G., Li, J. W., Genshaft, A. S., Kazer, S. W., Payer, K. R., de Riba Borrajo, J., Blainey, P. C., Irvine, D. J., Shalek, A. K. & Manalis, S. R. A microfluidic platform enabling singlecell RNA-seq of multigenerational lineages. Nat Commun 7, 10220 (2016).

6. Phillips, N. E., Mandic, A., Omidi, S., Naef, F. & Suter, D. M. Memory and relatedness of transcriptional activity in mammalian cell lineages. Nature Communications 10, 1208. issn: 2041-1723 (2019).

7. Muramoto, T., Muller, I., Thomas, G., Melvin, A. & Chubb, J. R. Methylation of H3K4 Is required for inheritance of active transcriptional states. Curr Biol 20, 397–406. issn: 1879-0445 0960-9822 (2010).

8. Shaffer, S. M., Emert, B. L., Reyes Hueros, R. A., Cote, C., Harmange, G., Schaff, D. L., Sizemore, A. E., Gupte, R., Torre, E., Singh, A., Bassett, D. S. & Raj, A. Memory Sequencing Reveals Heritable Single-Cell Gene Expression Programs Associated with Distinct Cellular Behaviors. en. Cell 182, 947–959.e17. issn: 00928674 (Aug. 2020).

9. Chang, H. H., Hemberg, M., Barahona, M., Ingber, D. E. & Huang, S. Transcriptome-wide noise controls lineage choice in mammalian progenitor cells. en. Nature 453, 544–547. issn: 0028-0836, 1476-4687 (May 2008).

10. Kalmar, T., Lim, C., Hayward, P., Munoz-Descalzo, S., Nichols, J., Garcia-Ojalvo, J. & Martinez Arias, A. Regulated fluctuations in nanog expression mediate cell fate decisions in embryonic stem cells. PLoS Biol 7, e1000149. issn: 1545-7885 (2009).

11. Hu, M., Krause, D., Greaves, M., Sharkis, S., Dexter, M., Heyworth, C. & Enver, T. Multilineage gene expression precedes commitment in the hemopoietic system. Genes & Development 11, 774–785. issn: 08909369 (Mar. 15, 1997).

12. Pina, C., Fugazza, C., Tipping, A. J., Brown, J., Soneji, S., Teles, J., Peterson, C. & Enver, T. Inferring rules of lineage commitment in haematopoiesis. Nature Cell Biology 14, 287–294. issn: 1465-7392, 1476-4679 (Mar. 2012).

13. Mojtahedi, M., Skupin, A., Zhou, J., Castañ, I. G., Leong-Quong, R. Y. Y., Chang, H., Trachana, K., Giuliani, A. & Huang, S. Cell Fate Decision as High-Dimensional Critical State Transition. en. PLOS Biology 14. Number: 12, e2000640. issn: 1545-7885 (Dec. 2016).

14. Richard, A., Boullu, L., Herbach, U., Bonnafoux, A., Morin, V., Vallin, E., Guillemin, A., Papili Gao, N., Gunawan, R., Cosette, J., Arnaud, O., Kupiec, J.-J., Espinasse, T., Gonin-Giraud, S. & Gandrillon, O. Single-Cell-Based Analysis Highlights a Surge in Cell-to-Cell Molecular Variability Preceding Irreversible Commitment in a Differentiation Process. en. PLOS Biology 14 (ed Teichmann, S. A.) e1002585. issn: 1545-7885 (Dec. 2016).

15. Moussy, A., Cosette, J., Parmentier, R., da Silva, C., Corre, G., Richard, A., Gandrillon, O., Stockholm, D. & Páldi, A. Integrated time-lapse and single-cell transcription studies highlight the variable and dynamic nature of human hematopoietic cell fate commitment. en. PLOS Biology 15 (ed Huang, S.) e2001867. issn: 1545-7885 (July 2017).

16. Gao, M., Ling, M., Tang, X., Wang, S., Xiao, X., Qiao, Y., Yang, W. & Yu, R. Comparison of High-Throughput Single-Cell RNA Sequencing Data Processing Pipelines en. preprint (Feb. 2020).

17. Moris, N., Edri, S., Seyres, D., Kulkarni, R., Domingues, A. F., Balayo, T., Frontini, M. & Pina, C. Histone Acetyltransferase KAT2A Stabilizes Pluripotency with Control of Transcriptional Heterogeneity. Stem Cells 36, 1828–1838. issn: 1066-5099 1066-5099 (2018).

18. Guillemin, A., Duchesne, R., Crauste, F., Gonin-Giraud, S. & Gandrillon, O. Drugs modulating stochastic gene expression affect the erythroid differentiation process. PLOS ONE 14, e0225166 (2019).

19. Stumpf, P. S., Smith, R. C. G., Lenz, M., Schuppert, A., Müller, F.-J., Babtie, A., Chan, T. E., Stumpf, M. P., Please, C. P., Howison, S. D., Arai, F. & MacArthur, B. D. Stem Cell Differentiation as a Non-Markov Stochastic Process. Cell Systems 5, 268–282 (2017).

20. Dussiau, C., Boussaroque, A., Gaillard, M., Bravetti, C., Zaroili, L., Knosp, C., Friedrich, C., Asquier, P., Willems, L., Quint, L., Bouscary, D., Fontenay, M., Espinasse, T., Plesa, A., Sujobert, P., Gandrillon, O. & Kosmider, O. Hematopoietic differentiation is characterized by a transient peak of entropy at a single-cell level. BMC Biology 20, 60. issn: 1741-7007 (2022).

21. Toh, K., Saunders, D., Verd, B. & Steventon, B. Zebrafish Neuromesodermal Progenitors Undergo a Critical State Transition in vivo. bioRxiv, 2022.02.25.481986 (2022).

22. Parmentier, R., Moussy, A., Chantalat, S., Racine, L., Sudharshan, R., Papili Gao, N., Stockholm, D., Corre, G., Fourel, G., Deleuze, J., Gunawan, R. & Paldi, A. Global genome decompaction leads to stochastic activation of gene expression as a first step toward fate commitment in human hematopoietic stem cells. bioRxiv (2021).

23. Gandrillon, O., Schmidt, U., Beug, H. & Samarut, J. TGF-Beta cooperates with TGF-Alpha to induce the self–renewal of normal erythrocytic progenitors: evidence for an autocrine mechanism. The EMBO Journal 18, 2764–2781. issn: 0261-4189, 1460-2075 (May 17, 1999).

24. Gandrillon, O. & Samarut, J. Role of the different RAR isoforms in controlling the erythrocytic differentiation sequence. Interference with the v-erbA and p135gag-myb-ets nuclear oncogenes. Oncogene 16, 563– 74 (1998).

25. Richard, A., Vallin, E., Romestaing, C., Roussel, D., Gandrillon, O. & Gonin-Giraud, S. Erythroid differentiation displays a peak of energy consumption concomitant with glycolytic metabolism rearrangements. PLoS One 14, e0221472. issn: 1932-6203 1932-6203 (2019).

26. Jaitin, D. A., Kenigsberg, E., Keren-Shaul, H., Elefant, N., Paul, F., Zaretsky, I., Mildner, A., Cohen, N., Jung, S., Tanay, A. & Amit, I. Massively Parallel Single-Cell RNA-Seq for Marker-Free Decomposition of Tissues into Cell Types. Science 343, 776–779. issn: 0036-8075, 1095-9203 (Feb. 14, 2014).

27. Woodworth, M. B., Girskis, K. M. & Walsh, C. A. Building a lineage from single cells: genetic techniques for cell lineage tracking. Nature Reviews Genetics 18, 230–244. issn: 1471-0056, 1471-0064 (Apr. 2017).

28. Zreika, S., Fourneaux, C., Vallin, E., Modolo, L., Seraphin, R., Moussy, A., Ventre, E., Bouvier, M., Ozier-Lafontaine, A., Bonnaffoux, A., Picard, F., Gandrillon, O. & Gonin-Giraud, S. Evidence for close molecular proximity between reverting and undifferentiated cells. BMC Biology 20, 155. issn: 1741-7007 (2022).

29. Terrén, I., Orrantia, A., Vitallé, J., Zenarruzabeitia, O. & Borrego, F. in Methods in Enzymology 239–255 (Elsevier, 2020). isbn: 978-0-12-818673-2.

30. Parish, C. R. Fluorescent dyes for lymphocyte migration and proliferation studies. Immunology and Cell Biology 77, 499–508. issn: 08189641 (Dec. 1999).

31. Kim, W., Klarmann, K. D. & Keller, J. R. Gfi-1 regulates the erythroid transcription factor network through Id2 repression in murine hematopoietic progenitor cells. Blood 124, 1586–1596 (Sept. 4, 2014).

32. Da Cunha, A. F., Brugnerotto, A. F., Duarte, A. d. S. S., Lanaro, C., Costa, G. G. L., Saad, S. T. O. & Costa, F. F. Global gene expression reveals a set of new genes involved in the modification of cells during erythroid differentiation: Modification of cells during erythroid differentiation. Cell Proliferation 43, 297–309. issn: 09607722, 13652184 (Apr. 28, 2010).

33. Aggarwal, C. C., Hinneburg, A. & Keim, D. A. On the Surprising Behavior of Distance Metrics in High Dimensional Space in Database Theory — ICDT 2001 (eds Van den Bussche, J. & Vianu, V.) (Springer Berlin Heidelberg, Berlin, Heidelberg, 2001), 420–434. isbn: 978-3-540-44503-

34. Benjamini, Y. & Hochberg, Y. Controlling the False Discovery Rate: A Practical and Powerful Approach to Multiple Testing. Journal of the Royal Statistical Society. Series B (Methodological) 57, 289–300. issn: 00359246 (1995).

35. Bonnaffoux, A., Herbach, U., Richard, A., Guillemin, A., Gonin-Giraud, S., Gros, P.-A. & Gandrillon, O. WASABI: a dynamic iterative framework for gene regulatory network inference. en. BMC Bioinformatics 20, 220. issn: 1471-2105 (Dec. 2019).

36. Wehling, A., Loeffler, D., Zhang, Y., Kull, T., Donato, C., Szczerba, B., Camargo Ortega, G., Lee, M., Moor, A., Gottgens, B., Aceto, N. & Schroeder, T. Combining single-cell tracking and omics improves blood stem cell fate regulator identification. Blood 140, 1482–1495. issn: 1528-0020 0006-4971 (2022).

37. Wang, F. & Higgins, J. M. Histone modifications and mitosis: countermarks, landmarks, and bookmarks. Trends Cell Biol 23, 175–84. issn: 1879-3088 0962-8924 (2013).

38. Golloshi, R., Sanders, J. T. & McCord, R. P. Genome organization during the cell cycle: unity in division. Wiley Interdiscip Rev Syst Biol Med 9. issn: 1939-005X 1939-005X (2017).

39. Palozola, K. C., Donahue, G. & Zaret, K. S. EU-RNA-seq for in vivo labeling and high throughput sequencing of nascent transcripts. STAR Protoc 2, 100651. issn: 2666-1667 2666-1667 (2021).

40. Kadauke, S., Udugama, M. I., Pawlicki, J. M., Achtman, J. C., Jain, D. P., Cheng, Y., Hardison, R. C. & Blobel, G. A. Tissue-specific mitotic bookmarking by hematopoietic transcription factor GATA1. Cell 150, 725–37. issn: 1097-4172 0092-8674 (2012).

41. Suter, D. M., Molina, N., Gatfield, D., Schneider, K., Schibler, U. & Naef, F. Mammalian genes are transcribed with widely different bursting kinetics. Science 332, 472–4. issn: 1095-9203 0036-8075 (2011).

42. Tunnacliffe, E. & Chubb, J. R. What Is a Transcriptional Burst? Trends Genet 36, 288–297. issn: 0168-9525 0168-9525 (2020).

43. Rodriguez, J. & Larson, D. R. Transcription in Living Cells: Molecular Mechanisms of Bursting. Annu Rev Biochem 89, 189–212. issn: 15454509 0066-4154 (2020).

44. Pedraza, J. M. & van Oudenaarden, A. Noise propagation in gene networks. Science 307, 1965–9 (2005).

45. Kim, S. & Shendure, J. Mechanisms of Interplay between Transcription Factors and the 3D Genome. Mol Cell 76, 306–319. issn: 1097-4164 1097-2765 (2019).

46. Martin-Martin, N., Carracedo, A. & Torrano, V. Metabolism and Transcription in Cancer: Merging Two Classic Tales. Front Cell Dev Biol 5, 119. issn: 2296-634X 2296-634X (2017).

47. Scrucca, L., Fop, M., Murphy, T. B. & Raftery, A. E. mclust 5: clustering, classification and density estimation using Gaussian finite mixture models. The R Journal 8, 289–317 (2016).

48. Di Tommaso, P., Chatzou, M., Floden, E. W., Barja, P. P., Palumbo, E. & Notredame, C. Nextflow enables reproducible computational workflows. Nature Biotechnology 35, 316–319. issn: 1087-0156, 1546-1696 (Apr. 2017).

49. R Core Team. R: A Language and Environment for Statistical Computing R Foundation for Statistical Computing (Vienna, Austria, 2021).

50. Cole, M. B., Risso, D., Wagner, A., DeTomaso, D., Ngai, J., Purdom, E., Dudoit, S. & Yosef, N. Performance Assessment and Selection of Normalization Procedures for Single-Cell RNA-Seq. Cell Systems 8, 315–328.e8. issn: 24054712 (Apr. 2019).

51. Breda, J., Zavolan, M. & van Nimwegen, E. Bayesian inference of gene expression states from single-cell RNA-seq data. Nature Biotechnology 39, 1008–1016. issn: 1087-0156, 1546-1696 (Aug. 2021).

52. Hafemeister, C. & Satija, R. Normalization and variance stabilization of single-cell RNA-seq data using regularized negative binomial regression. bioRxiv (Mar. 18, 2019).

53. Becht, E., McInnes, L., Healy, J., Dutertre, C.-A., Kwok, I. W. H., Ng, L. G., Ginhoux, F. & Newell, E. W. Dimensionality reduction for visualizing single-cell data using UMAP. Nature Biotechnology 37, 38– 44. issn: 1087-0156, 1546-1696 (Jan. 2019).

