## Supplementary file for "Differentiation is accompanied by a progressive loss in transcriptional memory"

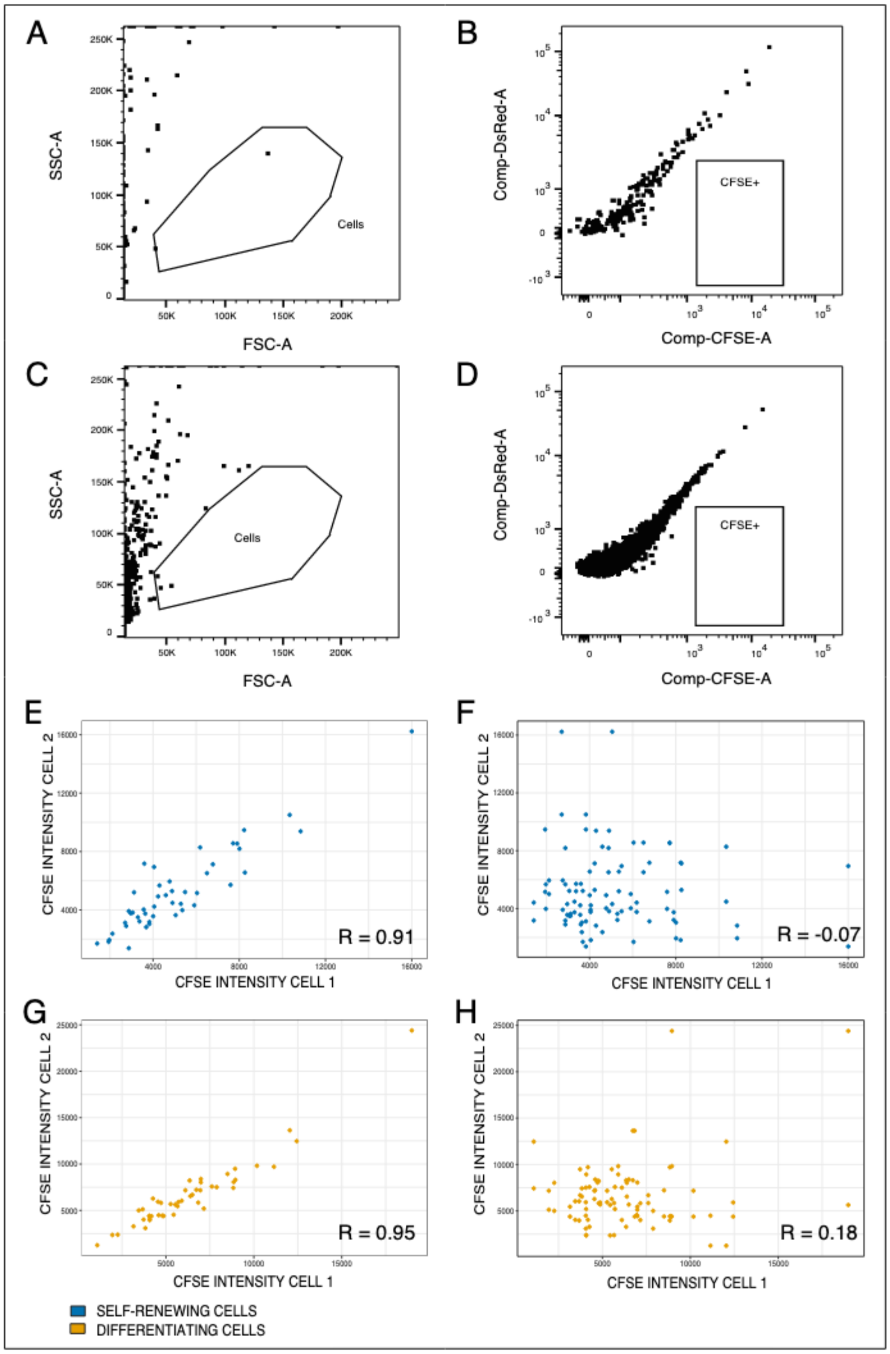


**Figure S1:** Technical validation of sister-cells isolation method using CFSE intensity data and evaluation of background noise for self-renewing and differentiating cells.

**Figure S1:** (A) Artefact detection in self-renewal medium. A few events are detected in the cell gate. (B) Artefact detection in self-renewal medium using CFSE signal (488nm, emission 530/30nm) versus auto-fluorescence (488nm, emission 585/42nm). No events is detected in the CFSE positive gate. For graphs A and B all events are displayed. (C) Artefact detection in differentiation medium. A few events are detected in the cell gate. (D) Artefact detection in differentiation medium using CFSE signal (488nm, emission 530/30nm) versus auto-fluorescence (488nm, emission 585/42nm). No events is detected in the CFSE positive gate. For graphs A and B all events are displayed. (E) Analysis of CFSE intensity correlation between self-renewing sister-cells (Spearman R = 0.91). (F) CFSE intensity correlation of randomly paired self-renewing cells (Spearman R = -0.07). (G) Analysis of CFSE intensity correlation between differentiating sister-cells (Spearman R = 0.95). (H) CFSE intensity correlation of randomly paired differentiating cells (Spearman R = 0.18).

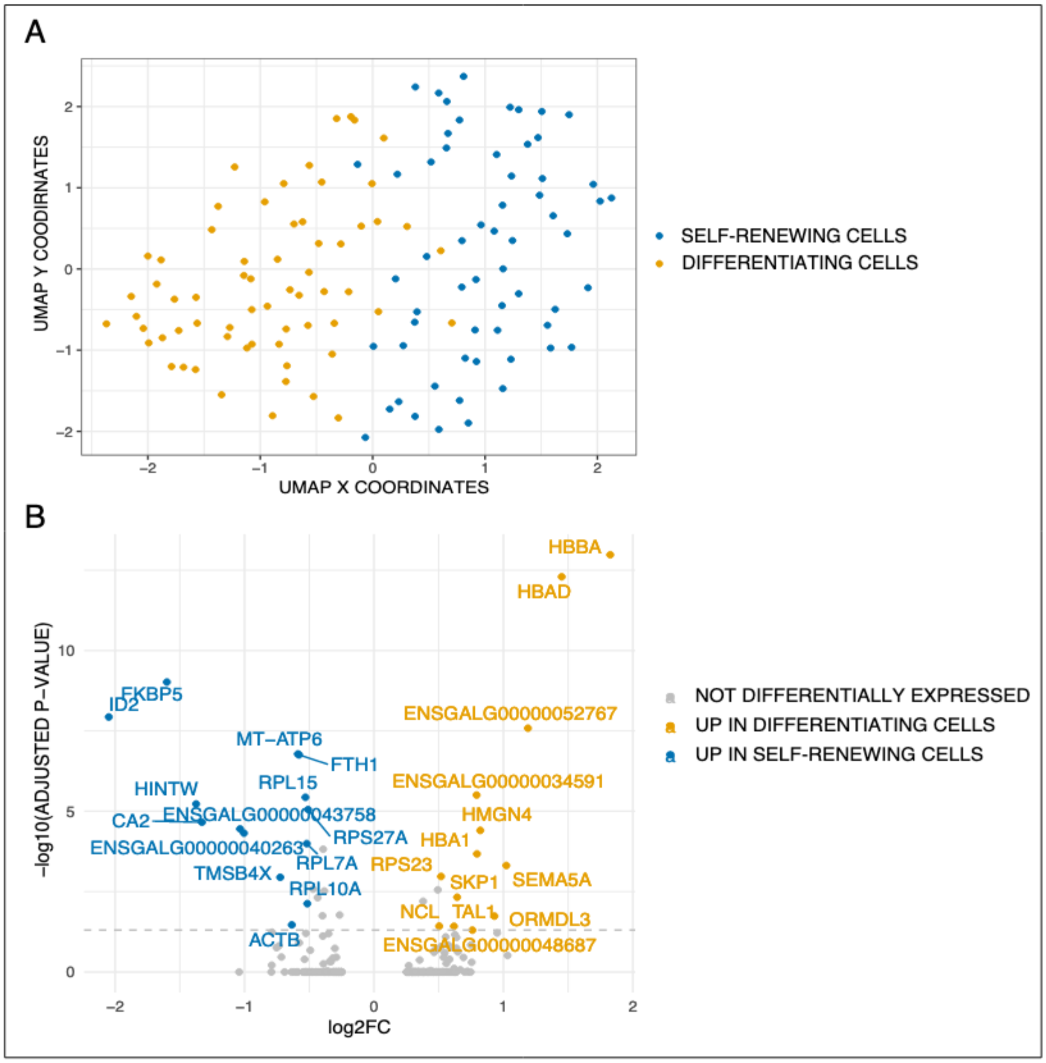


**Figure S2:** General structure of the data and characterization of the differentiation process.
(A) Dimensional reduction and projection with UMAP of the scRNA-seq data on T2EC cells. Cells in self-renewal are in blue and differentiating cells are in yellow. (B) Volcano plot of genes differentially expressed between the two conditions. Genes are considered significantly differentially expressed when the fold change is equal or above 0,5 and adjusted p-value is below 0.05 (grey dotted line). Blue dots represent significantly up-regulated genes in self-renewing cells and yellow dots represent significantly up-regulated genes in differentiating cells.

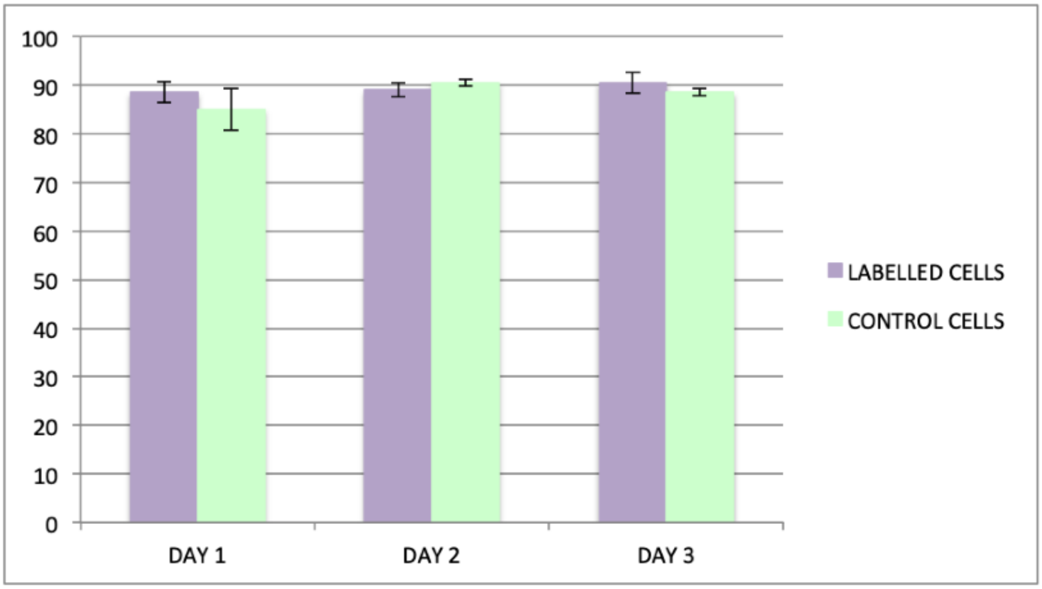


**Figure S3:** Histograms of cells viability.
Histograms of the percentage of viability, evaluated using Trypan Blue, during the cells staining (after day 1, 2 and 3) of the fluorescently labelled cells compared to negative control cells, on two biological replicates. T-test showed no significant differences.

| T2EC MEMORY GENES | |
| --- | --- |
| Gene_name | Ensembl_Gene_ID |
| ACTB | ENSGALG00000009621 |
| ATP5G3 | ENSGALG00000009286 |
| ATP6V0C | ENSGALG00000009229 |
| B2M | ENSGALG00000002160 |
| CCNG1 | ENSGALG00000001718 |
| CD99 | ENSGALG00000024488 |
| CLTA | ENSGALG00000015326 |
| DHRS7 | ENSGALG00000011921 |
| EEF1A1 | ENSGALG00000015917 |
| ENSGALG000 | ENSGALG00000040263 |
| ENSGALG000 | ENSGALG00000043758 |
| ENSGALG000 | ENSGALG00000050548 |
| ENSGALG000 | ENSGALG00000052767 |
| ENSGALG000 | ENSGALG00000053077 |
| ENSGALG000 | ENSGALG00000053765 |
| ESF1 | ENSGALG00000034768 |
| GAPDH | ENSGALG00000014442 |
| H2AFZ | ENSGALG00000014023 |
| HBA1 | ENSGALG00000043234 |
| HBAD | ENSGALG00000031597 |
| HBBA | ENSGALG00000047152 |
| HINTW | ENSGALG00000035998 |
| HMGB2 | ENSGALG00000010745 |
| HSP90AA1 | ENSGALG00000033212 |
| ID2 | ENSGALG00000035016 |
| KPNA2 | ENSGALG00000003584 |
| LBR | ENSGALG00000009305 |
| LDHA | ENSGALG00000006300 |
| LY6E | ENSGALG00000041621 |
| MLANA | ENSGALG00000019756 |
| MRPS28 | ENSGALG00000036749 |
| MT-ATP6 | ENSGALG00000041091 |
| MT-COX3 | ENSGALG00000035334 |
| MT-ND2 | ENSGALG00000043768 |
| PLK1 | ENSGALG00000006110 |
| PPIA | ENSGALG00000028600 |
| RHAG | ENSGALG00000016684 |
| RPL13 | ENSGALG00000006179 |
| RPL22L1 | ENSGALG00000009312 |
| RPL37 | ENSGALG00000014833 |
| RPS23 | ENSGALG00000015617 |
| RTFDC1 | ENSGALG00000007709 |
| SAT1 | ENSGALG00000016348 |
| SEMA5A | ENSGALG00000028685 |
| SH3BGRL3 | ENSGALG00000038536 |
| SMC2 | ENSGALG00000015691 |
| SOD1 | ENSGALG00000015844 |
| SPARC | ENSGALG00000004184 |
| ST13P5 | ENSGALG00000012007 |
| TFRC | ENSGALG00000007485 |
| TPD52 | ENSGALG00000040167 |
| TPX2 | ENSGALG00000006267 |
| TUBA1B | ENSGALG00000052192 |
| UBA52 | ENSGALG00000037716 |
| UBE2I | ENSGALG00000006428 |

| CD34+ MEMORY GENES | |
| --- | --- |
| Gene_name | Ensembl_Gene_ID |
| BCAT1 | ENSG00000060982 |
| GATA1 | ENSG00000102145 |
| HK1 | ENSG00000156515 |
| ACTB | ENSG00000075624 |
| KIT | ENSG00000157404 |
| CD38 | ENSG00000004468 |
| C22orf28 | ENSG00000100220 |
| ERG | ENSG00000157554 |
| CD133 | ENSG00000007062 |
| CD74 | ENSG00000019582 |

**Table S1:** Memory genes list with gene names and ENSEMBL gene ID, identified using linear models approach.
